# Diet outweighs genetics in shaping gut microbiomes in Asian honeybee

**DOI:** 10.1101/2022.01.23.477436

**Authors:** Qinzhi Su, Min Tang, Jiahui Hu, Junbo Tang, Xue Zhang, Xingan Li, Qingsheng Niu, Xuguo Zhou, Shiqi Luo, Xin Zhou

**Affiliations:** College of Food Science and Nutritional Engineering, China Agricultural University, Beijing, People’s Republic of China; Department of Entomology, College of Plant Protection, China Agricultural University, Beijing, People’s Republic of China; Key Laboratory for Bee Genetics and Breeding, Jilin Provincial Institute of Apicultural Sciences, Jilin Province, People’s Republic of China; Department of Entomology, University of Kentucky, Lexington, KY, USA

**Keywords:** gut microbiota, co-adaptation, population variation, pollen, nectar, adaptation

## Abstract

**Background:** The gut microbiome is a crucial element that facilitates a host’s adaptation to a changing environment. Host-specificity often coincides with distinctions in gut microbes, suggesting a co-evolution of the holobionts. However, it is unclear how gut microbiota shared by a common host ancestor would co-diversify with hosts and eventually become distinct among sister hosts. In this context, understanding the evolutionary pathway of gut microbiomes of the same host species could provide insight on how holobionts co-adapt along environmental gradients. Specifically, we ask which factor, nature or nurture, i.e., genetics or diets, contributes more to the shaping of gut microbiome, along with host diversification and range expansion.

**Results:** We compared and analyzed the gut microbiomes of 99 Asian honeybees, *Apis cerana,* from genetically diverged populations covering 13 provinces across China. Bacterial composition varied significantly across populations at phylotype, sequence-discrete population (SDP), and strain levels, but with extensive overlaps, indicating the diversity of microbial community among *A. cerana* populations is driven by nestedness. Taken together, genetics exhibited tangential impacts, while pollen diets were significantly correlated with both the composition and function of gut microbiome. Core bacteria, *Gilliamella* and *Lactobacillus* Firm-5, showed antagonistic turnovers and contributed to the enrichment in carbohydrate transport and metabolism. By feeding and inoculation bioassays, we confirmed that the variations in pollen polysaccharide composition contributed to the trade-off of these core bacteria.

**Conclusions:** Progressive change, i.e., nestedness, is the foundation of gut microbiome evolution in the Asian honeybee. Such a transition during the co-diversification of gut microbiomes is shaped primarily by environmental factors, diets in general, pollen polysaccharide in particular.

## Background

The gut microbiome often serves as a critical component in host’s adaptation to a changing environment [1]. Phylogenetically distant hosts consuming distinct diets have typically diverged for a long evolutionary time. Thus, it is not surprising that these hosts are revealed with abrupt differentiation in gut microbiomes, such as in broad mammal lineages [2]. Increasing evidence also indicates that the gut microbiomes of closely related host species, such as honeybees (*Apis* spp.), are also often host-specific [3]. Such a characteristic association and the fact that host-specific symbionts could often increase the overall fitness of the host, are considered as strong evidence for holobiont co-evolution [4].

From an evolutionary point of view, such beneficial symbionts could reach the most advantage through vertical transmission across host generations. In this regard, diverse transmission mechanisms have been reported in varied hosts, mostly involving eggs or special structures hosting symbionts as the vessel, e.g., bacteriocytes in aphids [5]. Especially, animals with parental care behaviors or sociality are often proofed as highly efficient in transferring crucial microbes to their offspring, e.g., humans [6], primates [7], birds [8], social bees [9], termites [10]. In such cases, gut microbiomes are considered inheritable within species. In the meantime, microbes currently specific to each of the closely related host species have likely derived from common ancestors that were already symbiotic to the common ancestor of the extant hosts [3]. However, it remains unclear how gut microbiota shared by a common host ancestor would co-diversify with hosts and eventually become distinct among extant sister hosts. As both host genetics and environment could have affected the evolution of gut microbiomes [11,12], it remains a challenge to understand the roles of nature and nurture in shaping gut microbiomes in a study system that involves distantly related hosts. In this context, natural populations of the same host species provide a proper system to investigate how the holobionts co-adapt and change along environmental gradients to elucidate the evolutionary pathway of the symbiotic bacteria.

In particular, for widespread species found in a large geographic range, environmental heterogeneity is expected to influence their gut microbiota [13,14]. This is because geographic location of animal populations is linked with varied host genetics, local vegetation, and environmental microbe sources. Unfortunately, most relevant studies focusing on intraspecific variation of symbionts only compared gut microbes in a few distinct populations or at a fine spatial scale [15–17]. Studies of large geographic scale were mainly conducted in humans, which is subject to confounding factors related to civilization, e.g., lifestyles, hygiene, antibiotics usage, travel [18–21]. In this regard, we know little about how gut microbiota naturally evolve under environmental heterogeneity on large geographic scales.

Honeybees (*Apis* spp.) may serve as an ideal model to understand the evolutionary dynamics between host and gut microbiota, in a natural setting. In particular, both the western (*A. mellifera*) and Asian honeybees (*A. cerana*) are widely distributed across tropical and temperate climates, each with endemic populations adapted to local habitats as the result of evolutionary processes [22,23]. Studies based on *A. mellifera* have established the framework for honeybee gut microbiota, revealing their essential role in honeybee biology, such as facilitating pollen digestion [24,25], host development [26] and pathogen resistance [27,28]. Varied honeybee species share much of the core microbes at the phylotype level, but possess host-specific microbial communities [3,29], showing distinct strain diversities among hosts [30]. These core microbes have probably become part of the symbiont system in the common ancestor of all extent corbiculate bees (honeybees, bumble bees, stingless bees, and orchid bees) [3]. However, little is known about how these gut symbionts have evolved within their hosts and eventually become distinct across honeybees, while remaining essential for the survival of each holobiont in its local habitat.

Compared to the highly managed *A. mellifera,* the Asian honeybee *A. cerana* remain mostly semi-feral across its natural range, including much of the Eastern, Southern, and Southeastern Asia [31]. Our recent work on the evolution of mainland *A. cerana* revealed that multiple peripheral subspecies had radiated from a common central ancestral population and adapted independently to the changing floras in diverse habitats [23]. In this system, both host genetics and changing floras could have served as determining factors for the formation of local honeybee gut profiles. Here, we aim to understand the landscape of gut microbial diversity and function across geographic populations of *A. cerana*. In particular, we query whether host genetics or diets have contributed more prominently to the formation of gut microbiomes in natural honeybee populations.

## Results

### Bacterial composition significantly varied across Asian honeybee populations at multiple levels

A total of 99 nurse bees from 36 hives, representing 15 geographic populations covering 13 provinces across China were analyzed (Fig. 1a). For each population, ≥5 gut samples were sequenced from at least two hives to represent the diversity of each population (Table S1). SNPs derived from honeybee reads were used to construct a neighbor-joining tree (Fig. 1b), which confirmed the geographic origin of the sampled populations. This result was consistent with the reported genetic structure and geographic distribution of *A. cerana* populations [23], thereby excluding the possibility of colony translocation.

**Fig. 1.**
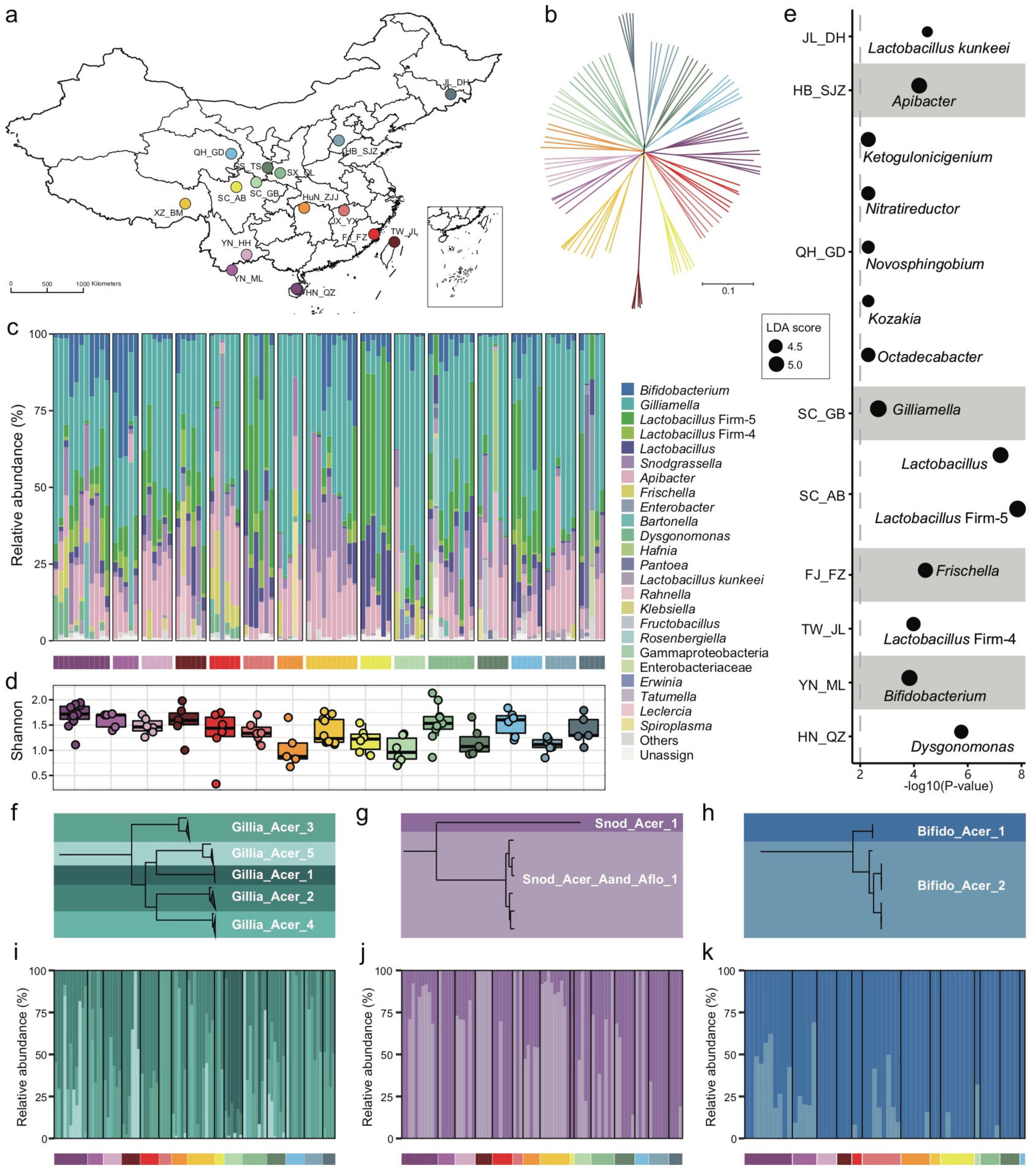
Bacterial composition of gut microbiota in geographic populations of *A. cerana*. (a) Sampling sites of 15 *A. cerana* geographic populations. (b) Neighbor-joining tree reflecting the honeybee population structure, based on genome-wide SNPs . Bacterial relative abundance (c) and Shannon index (d) based on gut metagenomes of different populations. Phylotypes with at least 5% abundance in any sample or 0.5% abundance in more than 6 samples were shown, otherwise included in “Others”. *Lactobacillus*: *Lactobacillus* that were not assigned to any known groups. (e) Featured gut microbe phylotypes in each geographic population revealed by LEfSe analyses. The size of the bubbles represents LDA score. Phylogenetic relationships of SDPs within *Gilliamella* (f), *Snodgrassella* (g) and *Bifidobacterium* (h). Maximal-likelihood phylograms, reconstructed using core genes present in all strains of the corresponding phylotype. The SDP compositions of *Gilliamella* (i), *Snodgrassella* (j) and *Bifidobacterium* (k) in gut samples, with those of abundances < 1% excluded. Horizontal bars under panels c and i-k indicate population origins of the guts, with colors corresponding to those in a and b.

Bacterial reads were then *de novo* assembled and aligned against the GenBank nr database to recover the phylotype composition for individual nurse bees. In congruence with previous studies [3,30], the core gut microbiota in *A. cerana* included six groups of bacteria, i.e. *Gilliamella* and *Snodgrassella* from Proteobacteria, *Bifidobacterium* from Actinobacteria, *Lactobacillus* Firm-4 and Firm-5 from Firmicutes, and *Apibacter* from Bacteroidetes (Fig. 1c). This result was further confirmed by the reference-based method (Figs. S1, S2), which employed a customized database containing 307 public and 83 newly sequenced bee gut bacterial genomes (Table S2). However, apparent variations in phylotype composition were observed among individual bees (Fig. 1c), and the composition of core-microbes appeared to be less stable than in *A. mellifera* [29,32,33].

Both Shannon index (Fig. 1d, Kruskal-Wallis, *P* = 0.0022) and phylotype diversity (ANOSIM, *r* = 0.29, *P* = 0.001) showed noticeable differences across populations of *A. cerana*. Nine of the fifteen investigated populations could be defined by featured bacteria in the LEfSe analysis [34], which showed significantly higher relative abundances in the focal population than all remaining populations (Fig. 1e).

The distinct gut variation across host populations was also reflected at finer taxonomic scales. Among all six core phylotypes in *A. cerana*, *Gilliamella* contained the most diverse host-specific sequence-discrete populations (SDPs) (Fig. 1f-h, Figs. S3-S6), which were defined as strains sharing a genome-wide average nucleotide identity (gANI) > 95% within each phylotype. Our results revealed varied presence and abundance in SDPs of core phylotypes among gut samples (Fig. 1i-k, Fig. S7), whereas *Gilliamella* showed significant SDP differences among geographical populations (Fig. S8, ANOSIM *r* = 0.14, *P* = 0.001). We also identified genome sites showing single nucleotide variation (SNV) for major SDPs in each sample, to detect gut variations at the strain level (Fig. S9). The results demonstrated significant variations in SNV composition across populations (Fig. S10). Thus, the gut bacterial composition of Asian honeybees varied significantly across geography at phylotype, SDP and strain levels.

### Progressive changes in honeybee microbial community were mainly determined by diets, not host genetics

Gut compositions showed extensive overlaps among populations, forming continuous groups in PCoA analyses (Fig. 2a), indicating progressive changes in microbial community structure among natural honeybee populations. Interestingly, a continuous variable contributing to the separation along the first principal coordinate axis (PCoA) reflected antagonistic dynamics in abundances of *Gilliamella* and *Lactobacillus* Firm-5 (Fig. 2b). Among all six core phylotypes, the relative abundance of *Gilliamella* (Spearman’s *rho* = -0.85, *P* = 2.14e-28) and *Lactobacillus* Firm-5 (Spearman’s *rho* = 0.79, *P* = 4.47e-22) showed the most significant correlation with the PCoA1 value.

**Fig. 2.**
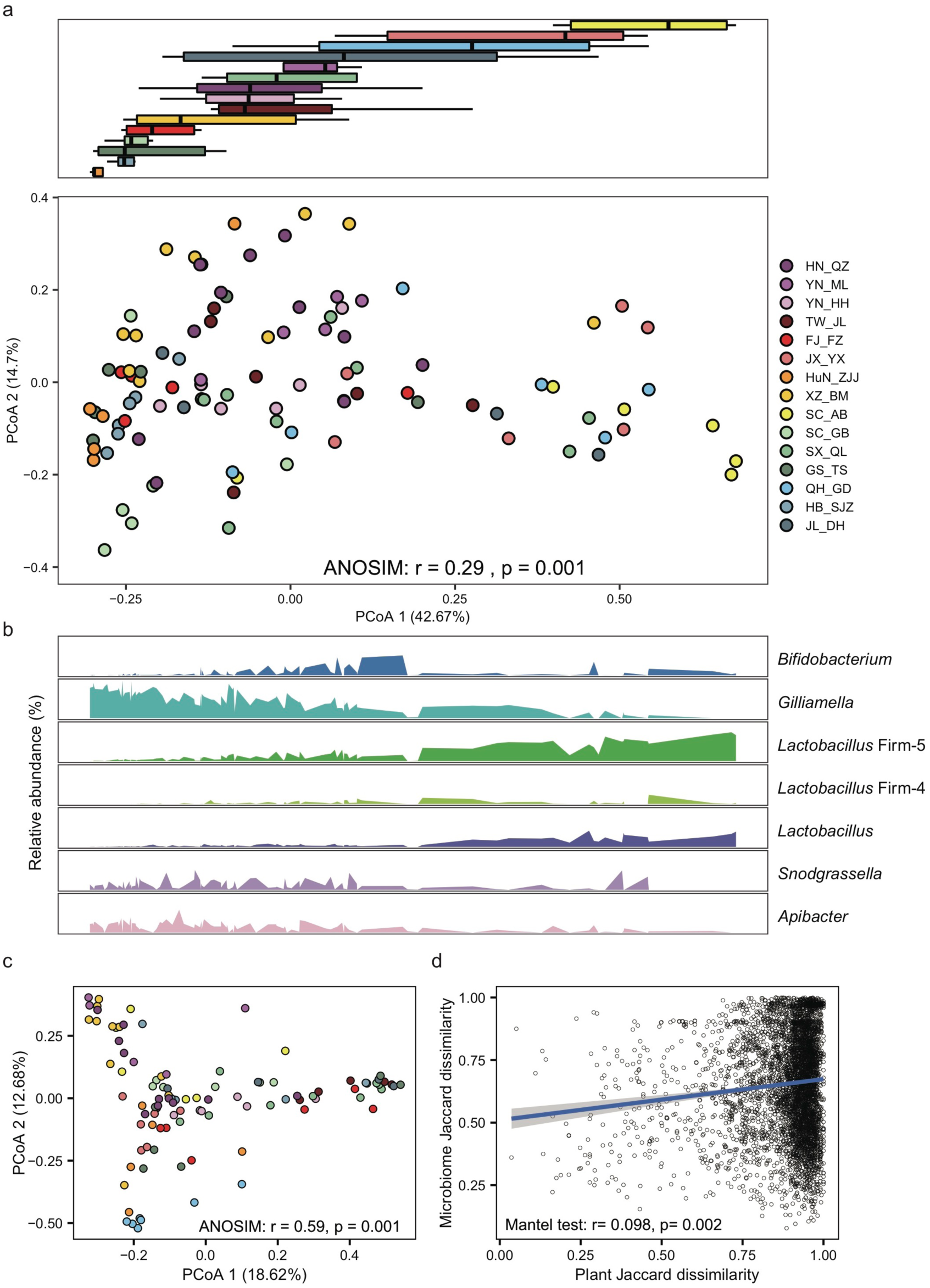
G*i*lliamella and *Lactobacillus* Firm-5 showed antagonistic trends in compositional turnover of honeybee gut microbiomes. (a) Overall variation of gut microbial community at the phylotype level, revealed by Bray-Curtis dissimilarity PCoA (bottom panel). Boxplots (top panel) indicate the distribution of each population along the first principal coordinate axis (PCoA1). Boxplot center values represent the median and error bars represent the SD. Colors correspond to the population origin of the gut samples. (b) Relative abundances of core bacterial phylotypes along PCoA1. (c) The pollen composition at the family level varied in gut metagenomes from populations of *A. cerana*. (d) The Jaccard distances of the gut bacterial phylotype and the pollen composition at the family level were significantly correlated.

To detect the impact of host genetics on gut microbiota, we estimated the heritability of the relative abundance of core bacteria at both phylotype and SDP levels. The heritability was overall low. Among the core phylotypes, *Gilliamella* abundance showed the highest heritability (Fig. S12), while that of *Snodgrassella* was not obvious. The abundances of about one third SDPs were not heritable. The GWAS analysis did not detect any apparent site variation that had determined bacterial composition, as no genomic region of *A. cerana* was found significantly associated with the bacterial abundance (with threshold as *P* < 2e-8) at either the phylotype or SDP level. These results indicated that gut microbial diversity at the geographic population level is not likely driven by host genetics, as measured by single-site nucleotide variations.

To examine the effect of diet on the gut microbiome, we first extracted pollen reads from the metagenome data and identified flower composition for each gut sample (details in Materials and Methods). Honeybee populations from different regions showed significant variation in pollen diet at the family level (ANOSIM, *r* = 0.59, *P* = 0.001, Fig. 2c, Table S3). Such a diet shift was further confirmed by pollen variation in honey samples extracted from five of the representative populations (SC_AB, SC_GB, SX_QL, QH_GD, JL_DH), where pollen composition again showed significant differences at the family level (ANOSIM, *r* = 0.35, *P* = 0.007, Fig. S11, Table S4). Most importantly, the Jaccard distances of the gut bacterial phylotype and the pollen composition were significantly correlated (mantel test, *r* = 0.098, *P* = 0.002, Fig. 2d). Among the core phylotypes, the abundances of *Gilliamella* showed significant correlation with the Shannon index of pollen composition in the gut (Spearman’s *rho* = -0.23, *P* = 0.020). Therefore, compared with host genetics, pollen diet showed predominant correlation with the composition of honeybee gut microbiome.

### KEGG Orthology (KO) function was correlated with diets and characterized in carbohydrate metabolism and transport

To understand whether gut microbes in *A. cerana* showed idiosyncratic regional traits on the function level, we *de novo* assembled the metagenomes and annotated genes for each of the 99 gut samples. As with bacterial compositions, the number of gene clusters per gut varied significantly among populations (Kruskal-Wallis test, *P*= 6.2e-4) (Fig. 3a). The gene cluster number in different individuals was significantly correlated with the Shannon index of gut bacteria (Pearson’s *r* = 0.64, *P* = 8.28e-13), suggesting that bacterial diversity is the basis for gene varieties among individual bees. We also quantified the rate of novel gene accumulation for each population. The results demonstrated distinct differences in gene diversity among populations (Fig. 3b).

**Fig. 3.**
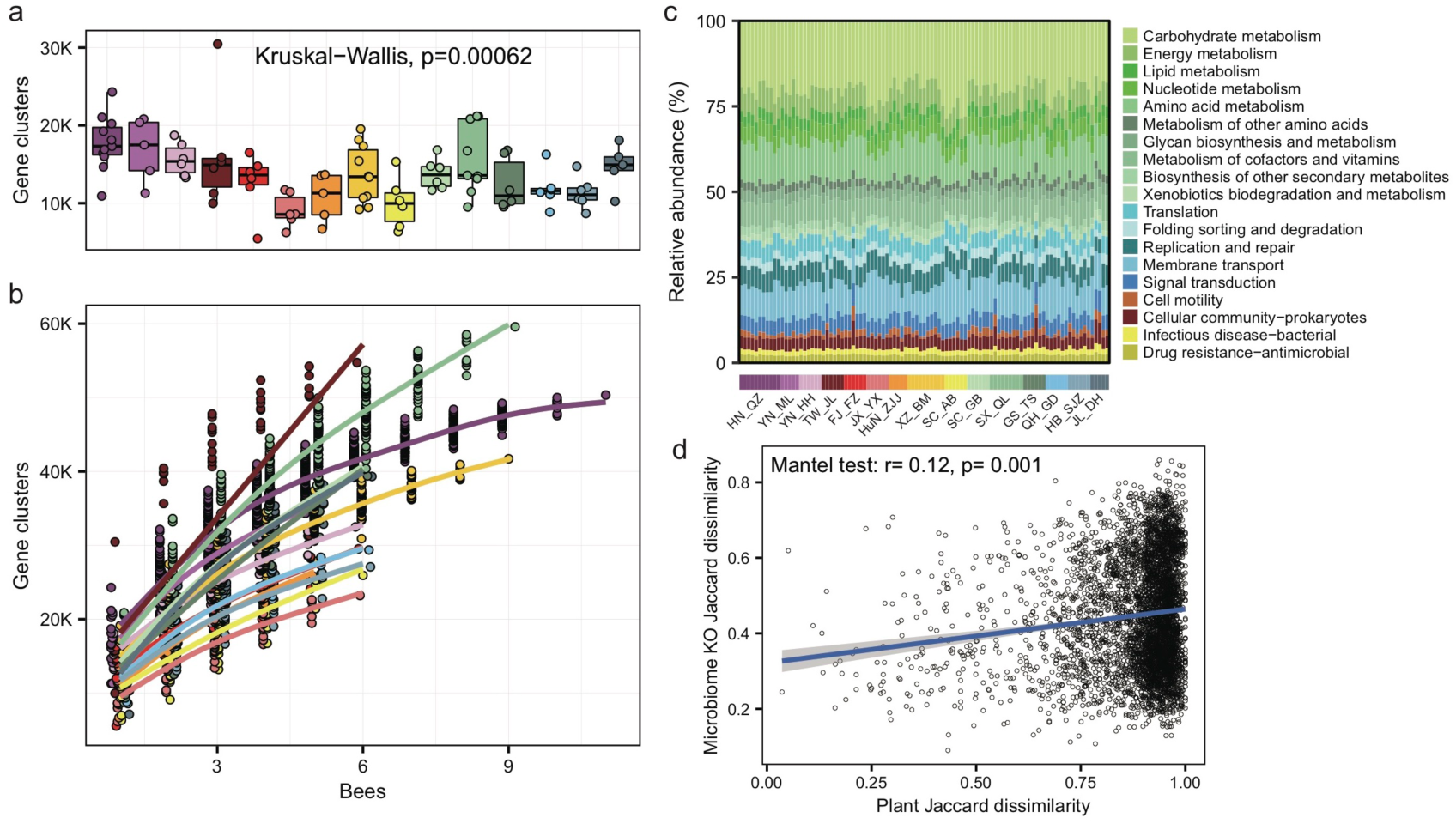
Significant variations in gene cluster and functional annotation among populations. (a) Gene cluster numbers per gut sample, based on 400 Mb bacteria-mapped reads. (b) Accumulation curves for gene clusters of each population of *A. cerana*, based on 400 Mb bacteria-mapped reads. (c) Relative abundance of KEGG annotations in each gut sample, based on all bacterium-mapped reads in metagenomes. (d) The Jaccard distances of the gut bacterial KO composition and the pollen composition at the family level were significantly correlated.

We assigned predicted gene clusters from all metagenome data to the KEGG database to reveal the diversity of functions among populations. A total of 1,965 functional orthologs (KOs) shared among all populations were enriched in genetic information processing, as well as signaling and cellular processes (Fig. S13). The KO category compositions (Fig. 3c) also showed extensive overlap, and were distinctively differentiated among populations (ANOSIM, *r* = 0.33, *P* = 0.001, Fig. S14). The LEfSe analyses showed that 11 of the 15 geographic populations had noticeably enriched KO categories (Fig. S15), which showed significantly higher relative abundances in the focal population than all remaining populations. The top significant population-enriched KOs (*P* < 1e-5) mainly included functions in metabolism and membrane transport (Fig. S14). Furthermore, the Jaccard distances of the gut bacterial KO composition and pollen diversity at the family level showed significant correlation (mantel test, *r* = 0.12, *P* = 0.001, Fig. 3d), indicating that not only bacterial composition but also their functions were associated with diets.

At the KO term level, we identified 83 KO terms showing inter-population differences (Table S5), in which they were significantly more abundant in only one of the geographic populations. Interestingly, 37 of 83 of the enriched KO terms were transporter pathway genes (all belonging to ko02000) (Fig. 4a), whereas the pathway was also enriched in some local populations (e.g., SC_AB and YN_ML, Fig. 4b). Most featured transporters were related to carbohydrates (Fig. 4c) and six of the enriched KO terms belonged to the glycoside hydrolase (GH) family (Table S5), in concert with the fact that polysaccharides are one of the major nutritional components derived from pollen. Therefore, the population-enriched gut microbe KOs were mainly associated with carbohydrate metabolism and transport, and were significantly correlated to pollen composition in a given local environment.

**Fig. 4.**
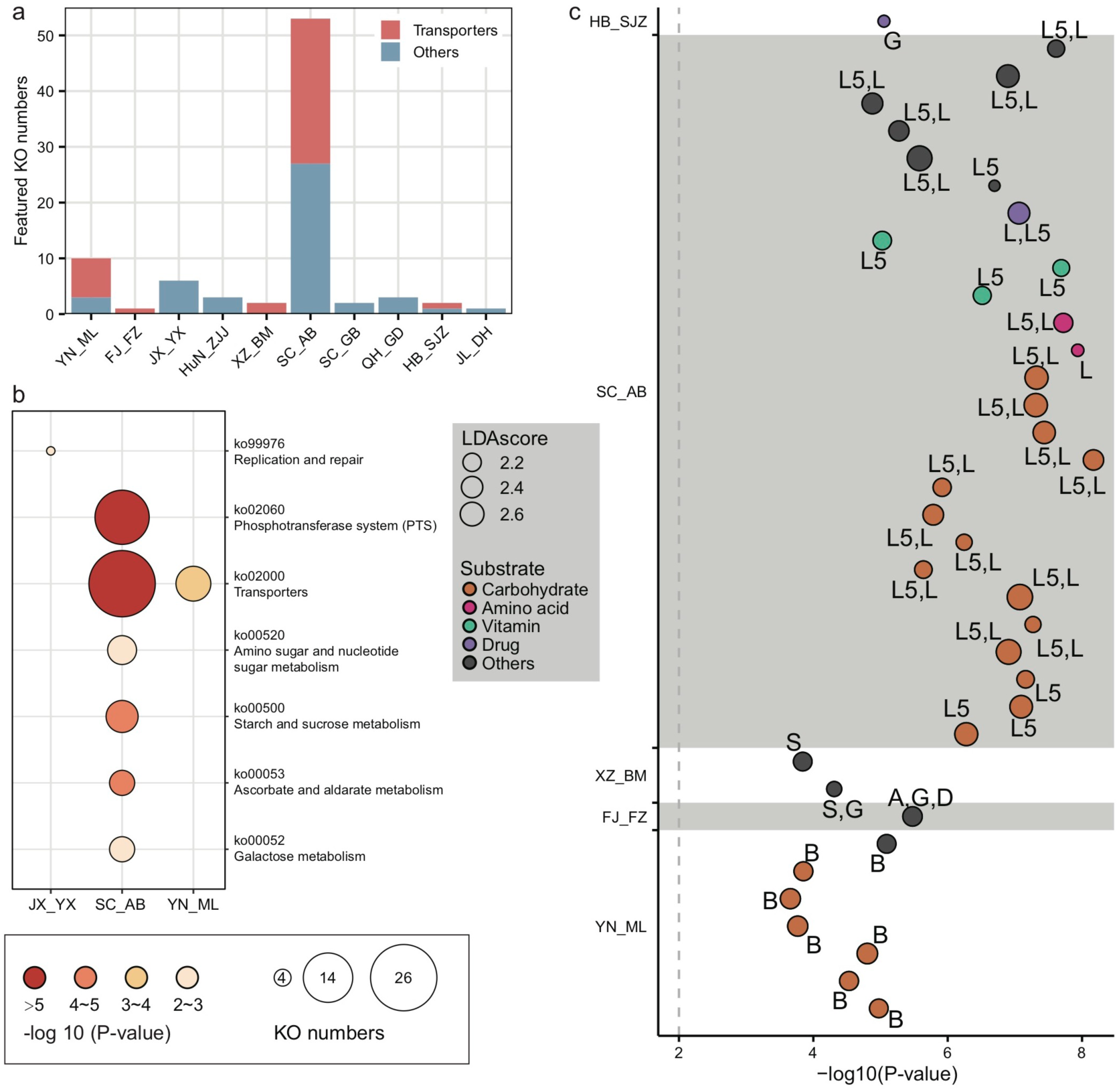
Locally featured KOs were enriched in carbohydrate transporters. (a) Featured KOs in geographic populations were enriched in transporters. (b) Featured KEGG pathways in gut microbiota from *A. cerana* populations. The size of the bubbles represents KO numbers. (c) Transporters in featured KOs were mainly specialized for carbohydrates. The size of the bubbles represents the LDA score. The codes marked next to each bubble indicate the main contributing bacteria species, where only those with > 10% contribution were listed: A: *Apibacter*; B: *Bifidobacterium*; D: *Dysgonomonas*; G: *Gilliamella*; L: *Lactobacillus* that is not assigned to any known groups; L5: *Lactobacillus* Firm-5; S: *Snodgrassella*.

### PTS, ABC transporters and GHs contributed by *Gilliamella*, *Lactobacillus* Firm-5 and *Bifidobacterium* were hotspot genes involved in local adaptation

In congruence with the finding that carbohydrate metabolism and transport play important roles in adapting to local diets, key genes of such pathways, such as phosphotransferase system (PTS) transporters and ATP binding cassette (ABC), were often characterized in distinct honeybee populations. For instance, a total of 17 PTS and 16 ABC transporters were identified from the 37 enriched transporter pathway genes (Table S5). All featured PTS genes were only found in the SC_AB population, while the featured ABC transporters were present in several populations (XZ_BM, SC_AB and YN_ML). PTS serves as one of the major mechanisms in carbohydrate uptake, particularly for hexoses and disaccharides. In SC_AB, the 17 featured PTS genes included some that are specific for ascorbate, beta-glucoside, cellobiose, fructoselysine/glucoselysine, galactitol, mannose, and sucrose (Table S5). The mapping of relevant gene clusters against the bacterial nr database suggested that these featured PTS genes were mainly contributed by *Gilliamella* and *Lactobacillus* Firm-5 (Table S6). The dominant role of these two bacteria in coding PTS genes was further confirmed by analyses of 81 individually sequenced and annotated genomes, where *Gilliamella* and *Lactobacillus* Firm-5 were the major phylotypes encoding PTS genes (Table S7). At the SDP level, *Lactobacillus* Firm-5 had a higher copy number of PTS transporters for cellobiose, fructoselysine/glucoselysine and galactitol than *Gilliamella* (Fig. 5a). Many of these PTS transporters were found in the featured genes in the SC_AB population, which was dominated by *Lactobacillus*. Thus, the enrichment of featured PTS genes could at least be partially explained by the elevated abundance of the contributing bacteria in local populations (Fig. 1e).

**Fig. 5.**
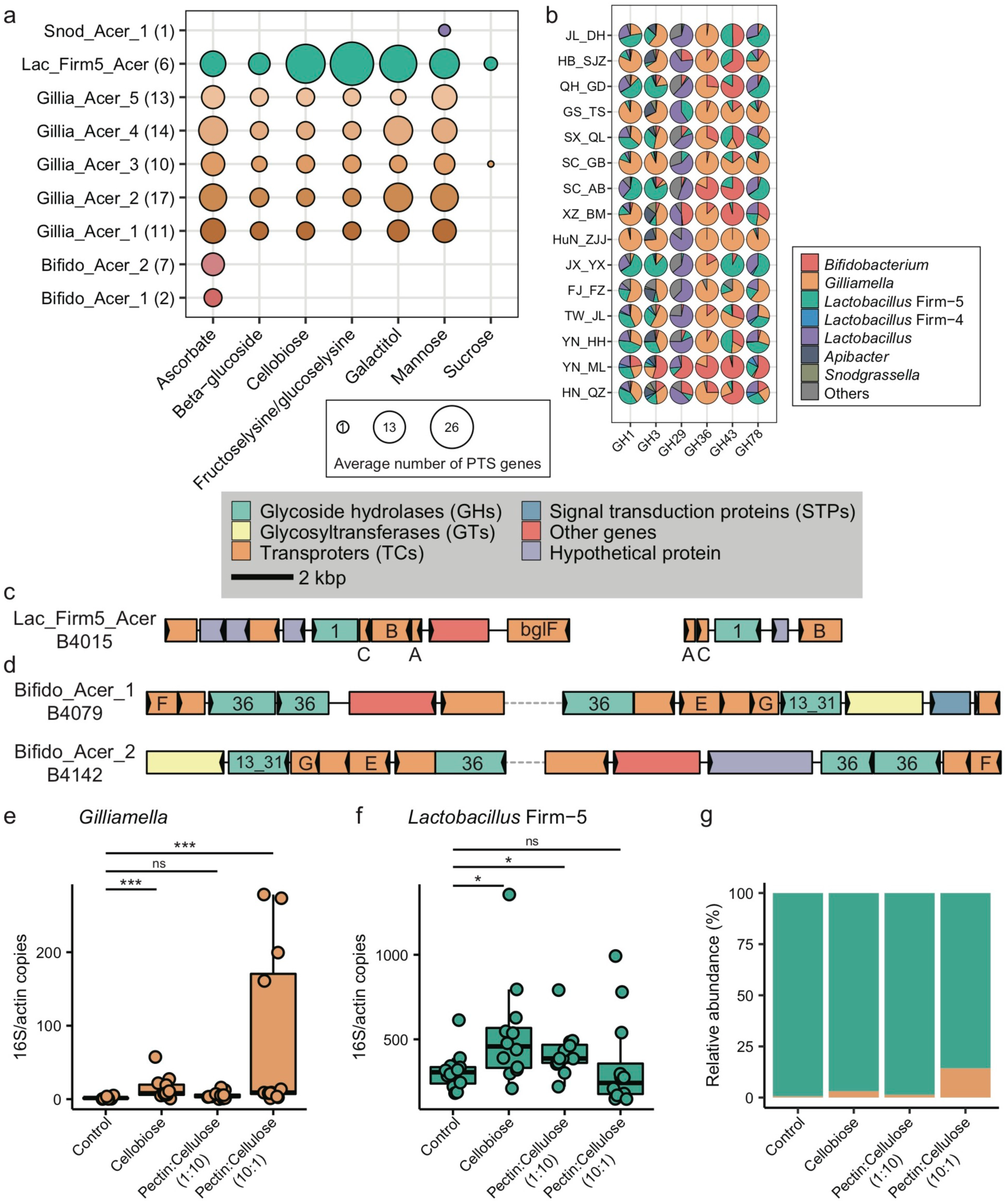
Main bacterial phylotypes coding for PTS and GHs. (a) Gene copy numbers in population-featured PTS pathways identified in all SDPs. Numbers in parentheses represent SDP strain numbers. Genes were annotated from the genomes of newly isolated microbial strains from *A. cerana* guts. (b) Featured GHs were coded by different bacterial phylotypes from metagenome of 15 geographic populations of *A. cerana*. (c) PTS transporters (*celA*/*celB*/*celC*/*bglF*), 6-phospho-beta-glucosidase (*bglA*) from the GH1 family were often found located together in genomes, which were represented here by *Lactobacillus* Firm-5 SDP. (d) ABC transporters (*msmE*/*msmF*/*msmG*), alpha-galactosidase from the GH36 family, and alpha-glucosidase from the GH13_31 family were often found located together in genomes, which were represented here by two *Bifidobacterium* SDPs. The change the absolute abundance of *Gilliamella* (e), *Lactobacillus* Firm-5 (f) and the percentage of *Gilliamella* and *Lactobacillus* Firm-5 (g) after feeding cellobiose and mixtures of pectin and cellulose with different concentrations. PTS: phosphotransferase system; GH: glycoside hydrolase. ABC: ATP binding cassette. ns: not significantly different, * *P* < 0.05, ** *P* < 0.01, *** *P* < 0.001.

The featured ABC transporters included transporters for amino acids, iron and carbohydrates (Table S5). Besides *Gilliamella* and *Lactobacillus* Firm-5, *Bifidobacterium* also contributed unique ABC transporters (Table S6). For example, the *Bifidobacterium*-unique transporters for raffinose/stachyose/melibiose (msmE, msmF and msmG) (genome annotation results in Table S7) were featured in the YN_ML population, in which *Bifidobacterium* was also the featured phylotype (Fig. 1e). The elevated *Bifidobacterium* and its unique ABC transporters characterized in YN_ML might be attributed to the presence of raffinose and stachyose in specific pollen or nectar, which are toxic to the honeybees [35].

At a finer taxonomic scale, 14 of the 17 featured PTS genes had significant population-distinct SNV sites coded by SDP from *Lactobacillus* Firm-5, and 9 of the 16 ABC transporters harbored significant population-distinct SNV sites coded by SDPs from *Lactobacillus* Firm-5 and *Apibacter* (Table S8). One featured gene *ulaC* (ascorbate PTS system EIIA or EIIAB component, K02821), coded by SDP from *Lactobacillus* Firm-5, showed significant population-distinct copy number variations (CNVs) (Table S9). Thus, the variations in functional genes seemed to have been caused by changes in the featured bacterial composition at both phylotype and strain levels.

Besides PTS and ABC genes, six GH genes were featured in *A. cerana* populations (from GH1, GH3, GH29, GH36, GH43 and GH78 family), and were mainly contributed by *Gilliamella*, *Lactobacillus* and *Bifidobacterium* (Fig. 5b, Table S10). To construct the profile for major gene families involved in glycoside breakdown, we used dbCAN2[36] to annotate all GH and polysaccharide lyase (PL) genes. We discovered that the GH/PL profiles varied across populations (Fig. S16). Additionally, non-core bacterium also encoded for novel GH genes. For instance, *Dysgonomonas* contributed unique GH gene families in *A. cerana*, including GH57, GH92, GH133 and GH144 (Table S10). This non-core bacterium was featured in the HN_QZ population (Fig. 1e), likely due to its contribution of unique GH gene sets.

Some of the six featured GH genes were positioned together with featured PTS or ABC transporters on the genome. Together, these genes formed CAZyme gene clusters (CGCs), performing sequential functions in polysaccharide degradation and transportation. For example, in *Lactobacillus* Firm-5, the featured 6-phospho-beta-glucosidase (*bglA*) from the GH1 family, PTS system genes for beta-glucoside and cellobiose were usually clustered and formed CGCs (Fig. 5c), and all these genes were enriched in the SC_AB population. In *Bifidobacterium*, the raffinose/stachyose/melibiose transport system msmEFG, and alpha-galactosidase from the GH36 family involved in raffinose/melibiose degradation were usually located together (Fig. 5d). These genes were all featured in the YN_ML population, which had *Bifidobacterium* as the featured phylotype.

### Feeding experiment verified the contribution of pollen polysaccharide composition to the trade-off of *Gilliamella* and *Lactobacillus* Firm-5

Our investigation on *A. cerana* guts from its natural range revealed antagonistic abundance between the two core-bacteria *Gilliamella* and *Lactobacillus* Firm-5 across geographic populations (Fig. 2b). As both lineage and function diversities of honeybee gut bacteria were strongly correlated to pollen diets (Figs. 2d, 3d), we speculate that characteristic traits in local food resources may have led to bacterial community shifts observed at the grand scale. To test this hypothesis, we conducted feeding experiments to verify whether functional adaptations observed in metagenomes can lead to adaptive advantages in bacterial competition.

As the main structural components of the pollen wall, pectin and cellulose were chosen as representative nutritional contents to examine the impacts of food on the abundance variation between *Gilliamella* and *Lactobacillus* Firm-5 in co-feeding experiments. In the main gut microbe phylotypes in honeybee, only *Gilliamella* are able to degrade the polygalacturonic acid (PGA), the backbone of pectin [3]. On the other hand, cellobiose (the key metabolite of cellulose) related PTS genes (Table S5) and metabolic pathway (ko00500, starch and sucrose metabolism) were highly enriched in the SC_AB population, as revealed by the metagenome data. The newly assembled *Lactobacillus* Firm-5 genome also showed elevated copy numbers in cellobiose PTS (Fig. 5a). As such, we anticipated that local food with higher proportion of pectin would increase the fitness of *Gilliamella*, and food with higher proportion of cellulose would favor *Lactobacillus* Firm-5 in the community.

We fed *A. cerana* workers that were colonized with equal abundance of *Gilliamella* and *Lactobacillus* Firm-5, with cellobiose, pectin and cellulose mixture with different concentrations (1:10 and 10:1 respectively) and examined corresponding changes in bacterial composition after four days. Interestingly, the absolute abundance of *Lactobacillus* Firm-5 was always higher than *Gilliamella* in the control group, which was only fed with sucrose (Fig. 5e-g), indicating a predominant role of *Lactobacillus* over *Gilliamella* in the given condition. The absence of pollen in food, and the absence of sucrose PTS genes in the strain we used (belonging to Gillia_Acer_2 SDP, Fig. 5a) might explain the low abundance of *Gilliamella* in the control group.

After feeding cellobiose, the absolute abundance of both *Gilliamella* and *Lactobacillus* Firm-5 significantly increased relative to the control group (Fig. 5e-f), which was in accordance with the presence of cellobiose PTS genes in both phylotypes (Fig. 5a). As expected, *Gilliamella* and *Lactobacillus* Firm-5 showed different responses to the mixed food with varied concentrations of pectin and cellulose. The absolute abundance of *Gilliamella* did not show significant variation after feeding food of pectin:cellulose (1:10), but the abundance of *Lactobacillus* Firm-5 significantly increased (Fig. 5e-f). On the other hand, the absolute abundance of *Gilliamella* showed significant increase after feeding food of pectin:cellulose (10:1), but the abundance of *Lactobacillus* Firm-5 did not vary significantly (Fig. 5e-f). The varied proportion of pectin and cellulose impacted the antagonistics of *Gilliamella* and *Lactobacillus* Firm-5. These results suggested that diet, pollen polysaccharide in particular, was the main driver in shaping gut bacterial composition and functions in *A. cerana*.

## Discussion

### Progressive change is the basis of gut microbiome evolution under local diet shift

In this study, we carried out comprehensive investigations on the gut microbiomes of the widespread Asian honeybee *A. cerana* at the population level. While many studies have contributed to our knowledge of the honeybee gut microbiota, little is understood about how this essential symbiont system is affected by changing environments and how it evolves with the host. In agreement with previous studies on both *A. mellifera* [33] and *A. cerana* [29], our results revealed variations in gut microbes among *A. cerana* individuals, even among those from the same hive. This individual distinction is expected, as worker bees obtain their gut microbiomes through social interactions [37], which is essentially a procedure of random subsampling from the bacterial pool maintained by cohabiting workers.

More importantly, our study revealed significant variations in gut microbiota across geographic populations of *A. cerana.* Our recent work on the evolution of mainland *A. cerana* revealed that selective pressures imposed by diverse habitats, especially those of the changing floras, led to convergent adaptation of the honeybee, where genes associated with sucrose sensitivity and foraging labor division had been repeatedly selected [23]. Here we showed that both microbial composition and function of the honeybee gut microbiota were highly dynamic throughout the studied natural range, along local adaptation of the honeybee hosts. Such an intra-species transition in gut microbiome reflects the evolutionary consequence of collective adaptation of both the honeybee and its symbiont.

In contrast to the abrupt distinction between *A. mellifera* and *A. cerana*, the gut microbiome of honeybee populations showed progressive change within host species (Fig. S17). Similarly, the gut microbiota community from 18 different human populations across geography also showed extensive overlap [38], implying a common trend for hosts exhibiting a continuous and wide-range distribution. Interestingly, changes in gut microbiome at the population level were closely correlated to the trade-off among core bacteria, both in honeybee and human. In our study, the two core bacteria *Gilliamella* and *Lactobacillus* Firm-5 showed antagonistic trends in occurrence in phylotype turnover across *A. cerana* populations. In the human guts, the trade-off of *Prevotellaceae* and *Bacteroidaceae* contributed to the first PC in PCoA of gut microbiota from different human populations in response to modernization [38]. The Russian population showed a different Bacteroidetes/Firmicutes ratio compared to the Western population [39].

### Factors shaping the gut microbiome: nature or nurture?

In humans, genome-wide association analysis identified some host factors in shaping microbiome [40–42]. However, in most cases, the lifestyle (e.g., foraging, traditional rural farming and urban industrialized life) over-dominates genetics [19]. Statistical analysis also demonstrated that environment dominated over host genetics in shaping human gut microbiota [43]. Similarly, the overwhelming role of nurture was also revealed in the Zucker rat, where age and local environment outweighed genetics in determining gut microbiome [44].

Among various environmental factors, the role of diet in human gut microbiome had been repeatedly addressed [45,46]. However, in natural human populations, diet seemed to have been frequently accompanied by other confounding factors related to lifestyles, such as culture, hygiene, and parasitic load. The complexity of human diet also made it difficult to identify precise dietary components and mechanisms that have modulated the gut microbiome. The honeybees, on the other hand, consume relatively simple but consistent food, i.e., pollen and honey, yet with variations in specific nutritional compositions. In this sense, the honeybees may serve as a better model to study how changes in nutrients would affect gut microbiota in a natural setting.

In our study, pollen analyses based on gut contents allowed us to establish strong associations between diet diversity and the Asian honeybee gut profiles, on both composition and function levels, while host genetics only exhibited tangential impacts. Diet with pollen was known to increase gut bacterial loads in Western honeybee [47]. Here, our feeding and inoculation assays further showed that pollen polysaccharide determined the abundance of the two core bacteria, *Gilliamella* and *Lactobacillus* Firm-5. The role of core-bacteria in local adaptations was reinforced by evidence showing their dominant contributions in genes related to pollen and nectar digestions. In particular, the PTS and ABC transporters, genes involved in the transportation of multiple types of polysaccharides, were primarily encoded by these two core bacteria, representing the most enriched transporters among all bacterial genes featured in local populations. We address that both PTS and ABC transporters were also highlighted in human populations from distant geographic regions. For example, the Russian population showed enriched PTS transporters compared to the Western population [39]. And the ABC transporters also showed enrichment in the rural population within Russia [39].

Unexpectedly, non-core bacteria sometimes became abundant in local honeybee populations. For instance, *Dysgonomonas* was typically low in abundance among *A. cerana* individuals, as reported in both *A. nigrocincta* [48] and *A. mellifera* [49]. But this bacterium contributed a series of unique GH genes in FJ_FZ and HN_QZ populations, thereby becoming abundant in the corresponding gut microbiome. This observation suggested that local food resources might trigger bacterial species turnover when non-core bacteria became more suited to new diets, which, again, highlighting the significance of diet on the gut profile.

### Population heterogeneity needs to be considered for the evolution and adaptation of honeybee microbiomes

A recent study suggested that both lineage and function diversities of the gut microbes were significantly lower in *A. cerana* when compared with *A. mellifera* [30]. However, as this conclusion was drawn based on two *A. mellifera* colonies from Switzerland, two colonies of both *A. mellifera* and *A. cerana* from two sites of Japan, it is difficult to anticipate whether such a distinct pattern could be generalized when population gradients of both honeybee species are taken into consideration. Although the present study was not designed for comprehensive analyses of inter-species comparisons, our results provided insights on how intra-species variations in gut microbiota might affect interpretations of differences between honeybee species.

Although the per-bee gene diversity was generally lower in *A. cerana* microbiota than *A. mellifera*, individual bees of several *A. cerana* populations (e.g., TW_JL, SX_QL) showed high levels of inter-individual variations (Fig. S18a). In addition, the divergence of the accumulated gene diversity between the two species was much less significant than previously suggested. The Japanese populations representing *A. cerana* in the earlier study [10] were one of the least variable populations among all *A. cerana* populations investigated in this study (Fig. S18b). Given the large variations observed among *A. cerana* populations, it is unknown whether a similar difference might also be common within *A. mellifera* and how that might influence the distinctions between these two widely distributed honeybee species. Additionally, other confounding factors should also be taken into consideration to gain a comprehensive understanding of honeybee gut microbiomes. In particular, the evolutionary pathways and phylogenetic relationships of focal populations, the specifics in honeybee management (such as colony merging and artificial diet additions) and other human interventions, may all have significant impacts on the honeybee gut profile. As the gut symbiont profile is a signature of natural adaptation of the holobiont to specific habitats, it would seem that comparisons between microbiomes of intra- and inter-host honeybee species should always be placed in a context of specific environments.

### A host-gut model that may help both honeybees and ourselves

As major agricultural pollinators, honeybees had experienced unprecedented global threats, such as the Colony Collapse Disorder [50], where corresponding changes in gut microbiomes were also noted [51]. Our study showed that the less-domesticated *A. cerana* had dynamic gut profiles corresponding to local diets. In this regard, regional floral diversity could serve as a key in maintaining characteristic repertoires of honeybee gut microbes, which is tremendously important for the honeybee health as a whole. Therefore, a sustaining plant community containing diverse endemic flower species should be considered as a key part of a honeybee conservation plan. On the other hand, the fitness of gut microbiomes of the honeybee populations may play an unforeseen role in the survival of colonies, during honeybee introduction, hybridization and especially translocation.

From a demographic perspective, understanding the honeybee gut system could also benefit human health. The honeybees passed their gut microbes through generations via social interactions, in a way that is very similar to the way humans do. Furthermore, the honeybee diet is confined to pollen and nectar, but diverse in nutritional combinations, providing an excellent opportunity for understanding dietary impacts on the formation of the gut symbiont system. Lastly, the divergence time among extant populations of *A. cerana* is relatively short, at ca. 100 ka [23], which is similar to that of the modern human populations [52–54]. Taken together, our analyses on gut microbiomes of *A. cerana* on the population level indicate that the honeybee is an ideal model to understand geographical variation of animal gut microbiota and the effect of diet on radiating populations.

## Conclusions

By sequencing the gut metagenomes and the genomes of isolated strains, we constructed the gut microbial profiles for *A. cerana* on both diversity and function levels. As the first attempt to characterize geographic dynamics of gut microbiota for natural honeybee populations, this study revealed that compositional and functional variations were common among geographic populations of *A. cerana*, both of which were significantly correlated to pollen composition recovered from gut metagenomes. The population variation in gut bacterial composition was closely correlated to *Gilliamella* and *Lactobacillus* Firm-5, which mainly contributed to population-featured functions in carbohydrate transport and metabolism. Our results uncovered the important roles of natural diet variation in shaping the gut microbiome in *A. cerana*. The results also add new insights into the progressive change of the gut microbe in a radiating species.

## Methods

### Sample collection

Nurse bees of *A. cerana* were obtained from inside the hives at 15 sites from 13 provinces of China (Hainan, Yunnan, Taiwan, Fujian, Jiangxi, Hunan, Tibet, Sichuan, Shannxi, Gansu, Qinghai, Hebei, and Jilin), between April 2017 and January 2019. Our sampling covers the main natural distribution range of *A. cerana* in China, from 19.2°-43.5°N, 95.7°-128.7°E, and from drastically different altitudes (12-3,325 m, Table S1). The guts (including the midgut and hindgut) were dissected from the abdomen and stored in 100% ethanol or directly frozen at -80 °C. To preserve live gut bacteria for strain isolation, a subset of guts was suspended in 100 μl of 25% glycerol (v/v, dissolved in PBS buffer), homogenized, and then frozen at -80 °C.

### Isolation, cultivation and identification of gut microbe strains

The gut homogenates were plated on different cultivation media respectively for various honeybee gut bacteria following Engel *et al*. [55], including heart infusion agar (HIA) with 5% (v/v) de-fibrinated sheep blood, Columbia agar with 5% (v/v) de-fibrinated sheep blood, De Man, Rogosa and Sharpe (MRS) agar, and trypticase-phytone-yeast (TPY) agar supplemented with 1% mupirocin. The plates were incubated at 35 °C in 5% CO_2_ or anaerobic atmosphere.

When bacterial colonies became visible on the plates, they were identified by sequences of their 16S rRNA gene. The isolates were picked and dissolved with H_2_O, then boiled at 100 °C for 1 min, which was used directly as DNA template in PCR. PCR amplicons were generated using the universal 16S primers 27F (5’-AGAGTTTGATCCTGGCTCAG-3’) and 1492R (5’-GGTTACCTTGTTACGACTT-3’) with 25 cycles of amplification (94 °C for 30 s, 60 °C for 40 s and 72 °C for 60 s) after an initial incubation for 1 min at 95 °C. Amplicons were sequenced using Sanger sequencing and identified using blastn against annotated sequences in GenBank.

### DNA extraction for genome and metagenome sequencing

The gut DNA was extracted following Kwong *et al.* [3]. Briefly, the crushed gut was suspended in a capped tube with 728 μl of CTAB buffer, 20 μl of proteinase K, 500 μl of 0.1-mm Zirconia beads (BioSpec), 2 μl of 2-Mercaptoethanol and 2 μl of RNase A cocktail. The mixtures were bead-beaten for 2 min for 3 times. After digested overnight at 50 °C, the mixtures were added with 750 μl of phenol/chloroform/isoamyl alcohol (25:24:1, pH 8.0) and centrifuged to obtain the aqueous layer. After being precipitated at -20 °C, spun at 4 °C and washed with -20 °C ethanol, the DNA pellets were dried at 50 °C and then re-suspended in 50 μl of nuclease-free H_2_O. Final DNA samples were stored at -20 °C.

Genomic DNA of honeybee gut bacterial isolates was also extracted using the phenol-chloroform protocol. The bacterial cells were re-suspended in 500 μl of lysis buffer [50 mM Tris-HCl (pH 8.0), 200 mM NaCl, 20 mM EDTA, nuclease-free H_2_O, 2% SDS, proteinase K (20 mg/ml)], then added with 500 μl of CTAB extraction buffer [50 mM Tris-HCl (pH 8.0), 20 mM EDTA, 1.4 M NaCl, 2% CTAB, 1% PVP 40000, nuclease-free H_2_O; pre-heated at 56 °C]. The mixtures were incubated for 30 min at 65 °C before the addition of 500 μl of phenol/chloroform/isoamyl alcohol (25:24:1, pH 8.0). Then the mixture was centrifuged at 14,000 g at room temperature (RT) for 5 min. The aqueous layer was transferred to a new tube, added with 5 μl of RNase (100 mg/ml), and incubated at RT for 20 min and added with 600 μl of chloroform: isoamyl alcohol (24:1). After spinning at 14,000 g at RT for 5 min, the aqueous layer was transferred to a new tube and added with 5 μl of ammonium acetate (final concentration 0.75 M), 1 μl glycogen solution (20 mg/ml) and 1 ml of cold 100% ethanol. DNA was precipitated at -20 °C for 30 min. Precipitations were spun at 14,000 g at 4 °C for 15 min, and the supernatant was decanted. DNA pellets were washed with 80% and 70% ethanol pre-cooled at -20 °C respectively and spun for an additional 10 min at 4 °C. The supernatant was discarded and the DNA pellet was air dried. The pellet was re-suspended in 50 μl nuclease-free H_2_O, and kept at 4 °C overnight before stored at -20 °C.

### Genome and metagenome sequencing

A total of 99 honeybee gut samples were used for metagenome sequencing (Table S1). And 83 isolated core bacterial strains obtained from *A. cerana* were also sequenced (Table S2). DNA samples were paired-end sequenced at BGI-Shenzhen using the BGISEQ-500 platform (200-400 bp insert size; 100 bp read length; paired-ended [PE]) and at Novogen using the Illumina Hiseq X Ten platform (350 bp insert size; 150 bp read length; PE). One *Gilliamella* strain (B3022) was sequenced on the PacBio RS II platform at NextOmics.

### Bacterial genome assembly and annotation

Low quality reads from the Illumina Hiseq X Ten platform were filtered out using fastp [56] (version 0.13.1, -q 20 -u 10) before subsequent analyses. For isolated bacterial strains, clean data were assembled using SOAPdenovo [57] (version 2.04, -K 51 -m 91 –R for PE 150 reads; -K 31 -m 63 –R for PE 100 reads), SOAPdenovo-Trans [58] (version 1.02, -K 81 -d 5 -t 1 -e 5 for PE 150 reads; -K 61 -d 5 -t 1 -e 5 for PE 100 reads), and SPAdes [59] (version 3.13.0, -k 33,55,77,85) based on contigs assembled by SOAPdenovo (only for PE 150 reads) or SOAPdenovo-Trans. The assembly with the longest N50 was retained for each strain as the draft genome. Then clean reads were mapped to the assembled scaftigs using minimap 2-2.9 [60] and the bam files were generated by samtools [61] (version 1.8). Genome assemblies were processed by BamDeal (https://github.com/BGI-shenzhen/BamDeal, version 0.19) to calculate and visualize the sequencing coverage and GC content of the assembled scaftigs. Scaftigs with aberrant depths and GC contents were then removed from the draft genome. Next, the remaining scaftigs were filtered taxonomically. Scaftigs assigned to eukaryote by Kraken2 [62] using the standard reference database were removed, and the ones aligned to a wrong phylum by blastn (megablast with e < 0.001) were further removed. The remaining genome assemblies were used as bacterial genome references. The *Giliamella* strain (B3022) sequenced on the PacBio RS II platform was assembled using a hierarchical genome assembly method (HGAP2.3.0) [63].

The protein coding regions of bacterial genomes were predicted using Prokka version 1.13 [64]. The KEGG orthologous groups (KOs) annotation was carried out using KofamKOALA [65] based on profile HMM and adaptive score threshold with default parameters. Programs KEGG Pathway and Brite Hierarchy were used to screen the annotation results. Finally, dbCAN2 version 2.0.11 [36] was applied to annotate CAZymes and CGCs using embedded tools HMMER, DIAMOND and Hotpep with default parameters.

### Genetic variation of *A. cerana* hosts

Metagenomes were filtered by fastp (-q 20 -u 10) [56]. Clean reads were then mapped to the *A. cerana* reference genome (ACSNU-2.0, GCF_001442555.1) [66] using the BWA-MEM algorithm (v 0.7.17-r1188) [67], with default settings and an additional “-M” parameter to reach compatibility with Picard. Read duplicates were marked using Picard MarkDuplicates 2.18.9 (http://broadinstitute.github.io/picard/). GATK HaplotypeCaller in the GVCF mode [68] (v4.0.4) was used to call variants for each sample. All of the per-sample GVCFs were joined using GenotypeGVCFs. Then the final variant file retained SNPs that met all of the following criteria: 1) average depth > 5× and < 40×; 2) quality score (QUAL) > 20; 3) average genotype quality (GQ) > 20; 4) minor allele frequency (MAF) > 0.05; 5) proportion of missing genotypes < 50%; 6) bi-allelic SNP sites.

The identity by state (IBS) distance matrices were performed and constructed with the filtered SNPs using functions “snpgdsIBS” in the R package SNPRelate [69]. A neighbor-joining tree was reconstructed based on the IBS distance matrix using the function “nj” in the R package Ape [70]. Node support values were obtained after 1,000 bootstrap replicates.

### Reference-based metagenome composition analyses

Shotgun reads generated from whole honeybee gut were firstly mapped against the *A. cerana* genome (ACSNU-2.0, GCF_001442555.1) using BWA aln (version 0.7.16a-r1181, -n 1) [67] to identify host reads, which were subsequently excluded. For taxonomic assignments of bacterial sequences, we used Kraken2 [62] and Bracken version 2.0 [71] to profile bacterial phylotype composition and used MIDAS [72] to profile strain composition for metagenomic samples. The reference database contained 390 bacterial genomes, including 307 published genomes and 83 newly-sequenced *A. cerana*-derived strains from this study (Table S2). The majority of the reference strains belonged to six core phylotypes (*Gilliamella, Snodgrassella, Bifidobacterium, Lactobacillus* Firm-4*, Lactobacillus* Firm-5, *Apibacter*) of honeybee gut bacteria. The analyses of public gut metagenome data of *A. cerana* from Japan [30] and *A. mellifera* [30,33] followed the same pipeline.

### Identification and profiling of SDP

We defined SDPs for each core gut bacterium (*Gilliamella*, *Snodgrassella*, *Bifidobacterium*, *Lactobacillus* Firm-4, *Lactobacillus* Firm-5, *Apibacter*) using a 95% gANI threshold [73]. Pairwise average nucleotide identities were calculated using the pyani Python3 module (https://github.com/widdowquinn/pyani). To generate the whole-genome tree for each core bacterium, we used Roary version 3.12.0 [74] with the parameter -blastp 75 to obtain core single-copy genes shared among all strains. The alignments of nucleotide sequences were concatenated, from which a maximum-likelihood tree was inferred using FastTree version 2.1.10 [75] with a generalized time-reversible (GTR) model and then visualized using iTOL [76].

We used the ‘run_midas.py species’ script of MIDAS [72] with default parameters to estimate SDP relative abundances in each sample. The script ‘merge_midas.py species’ with the option ‘--sample_depth 10.0’ was used to merge SDP abundance files across samples. The SDPs with a relative abundance less than 1% were filtered out.

### Detection of SNV and CNV across populations

CheckM version 1.0.86 [77] was used to estimate the completeness and contamination of genomes. The genome with the highest completeness and lowest contamination was selected as the reference sequence for each SDP. The metagenomic reads were mapped against reference genomes and the SNVs were quantified along the entire genome using MIDAS [72] and the script ‘run_midas.py snps’ with default parameters. For each SDP, the script ‘merge_midas.py snps’ pooled data across multiple samples with options ‘--snp_type bi --site_depth 5 --site_prev 0.05 – sample_depth 5.0 –fract_cov 0.4 –allele_freq 0.01’ to obtain the minor allele (second most common) frequency file. Thus, bi-allelic SNVs prevalent in more than 5% of profiled samples were predicted and rare SNVs with abnormally high read depth were excluded. The matrix files of SNVs remaining polymorphic were obtained after filtering steps.

We used the ‘run_midas.py genes’ script in MIDAS [72] to map metagenomic reads to pangenomes of each SDP and quantified gene copy numbers with default parameters. Then we merged results from pangenome profiling across samples with the option ‘--sample_depth 5.0’ from the ‘merge_midas.py genes’ module. The gene coverage was normalized by the coverage of a set of 15 universal marker genes to obtain the estimated copy number for genes of each SDP. The coverage of each KO term was obtained by summing up all genes annotated as the same KO for each SDP. *P* values were calculated using the Kruskal-Wallis one-way analysis across populations with the ‘compare_means’ function in the R package ‘ggpubr’. KO copy number variation and SNV of each SDP were detected as highly variable when an adjusted *P* value < 0.05.

### *De novo* assembly of metagenomes

The metagenome was also *de novo* assembled using MEGAHIT [78] (version 1.1.2, -m 0.6 --k-list 31,51,71 --no-mercy) for each gut sample. Assemblies longer than 500 bp were blasted against the NCBI nr database using DIAMOND [79] (version 0.9.22.123, blastx -f 102 -k 1 -e 1e-3) and were assigned to fungi, bacteria, archaea, virus or plants (Viridiplantae). Only assemblies assigned as bacteria were retained for further analyses.

A customized bacterial genome database was constructed to enable taxonomic assignments for the bacterial assemblies. The database included all bacterial genomes available on NCBI (ftp://ftp.ncbi.nlm.nih.gov/genomes/genbank/bacteria/) up to Jan 2019 (167,172 genomes), 83 genome assemblies of newly sequenced *A. cerena* gut bacteria (Table S2) and 14 *Apibacter* genomes from *A. cerana* [80]. Taxonomical assignments were conducted using blastn and an e-value of 1e-5. The assemblies were assigned to the genus of the best hit, while those without any hits were defined as unassigned bacteria.

For each metagenome sample, all clean reads were mapped against bacterial assemblies using SOAPaligner [81] (version 2.21, -M 4 -l 30 -r 1 -v 6 -m 200). The results were summarized using the soap.coverage script (version 2.7.7, http://soap.genomics.org.cn/down/soap.coverage.tar.gz). Only assemblies with ≥ 90% coverage were considered as true bacteria. Shannon index and Bray-Curtis dissimilarity were calculated using the vegan R package [82]. The analyses of public gut metagenome data of *A. cerana* from Japan [30] and *A. mellifera* [33] followed the same pipeline.

### Gene prediction and functional annotation for metagenomes

Gene prediction was conducted using MetaGeneMark [83] (GeneMark.hmm version 3.38) with the *de novo* metagenome assemblies, and those longer than 100 bp were clustered using CD-HIT [84] (version 4.7, -c 0.95 -G 0 -g 1 -aS 0.9 -M 0) to obtain a non-redundant gene catalog for *A. cerana* metagenomes. For each individual metagenome sample, clean data were aligned onto the non-redundant gene catalog using SOAPaligner [81] (version 2.21, -M 4 -l 30 -r 1 -v 6 -m 200). And the gene abundance was calculated using the soap.coverage script (version 2.7.7, http://soap.genomics.org.cn/down/soap.coverage.tar.gz). For each sample, only assemblies of ≥ 90% coverage were retained for further annotation. The analyses of public gut metagenome data of *A. cerana* from Japan [30] and *A. mellifera* [33] followed the same pipeline.

Functional annotation of gene catalog was performed by GhostKOALA [85] using the genus_prokaryotes KEGG GENES database and KofamKOALA [65] with an e-value threshold of 0.001. Genes were firstly assigned with KO ID predicted by KofamKOALA, and the remaining unassigned genes were then annotated using GhostKOALA. KOs were mapped onto KEGG pathways using the KEGG Mapper online (https://www.kegg.jp/kegg/tool/map_pathway2.html).

The abundances of KOs and pathways were calculated as the sum of the abundances of all genes annotated to them using custom scripts. Population dissimilarities (Bray–Curtis distance) of KO function among the 15 bee populations were tested by the ANOSIM test included in the vegan package [82] with 999 permutations. Linear discriminant analysis (LDA) was performed using LEfSe [34] with default parameters to identify KO biomarkers in different populations. Function enrichment of featured KOs was estimated by one-sided Fisher’s Exact Test using the stats R package at both module and pathway levels.

For each featured KO, the abundances for all bacterial species encoding the KO-related genes were listed for all of the 99 samples. In each population, the median abundance was used as the abundance of bacterial species encoding the respective KO. Then the contributions by different bacterial species to the corresponding KO were estimated. If the KO term was identified in > 50% individual bee guts of the same population, the KO was considered to be present in the population. To compare gene numbers among different populations, we standardized metagenome data by randomly extracting 400 Mb bacterium-derived data from each gut sample, which were mapped to the gene assemblies. The assemblies were retained only if the coverage ≥ 90%.

GH and PL genes were functionally assigned with the dbCAN2 database [36]. In each population, the median abundance was used as the abundance of bacterial species encoding respective GH/PL gene clusters. Then the contributions by different bacterial species to the corresponding GH/PL gene clusters were estimated.

### Diet profiling of gut and honey metagenomes

A customized chloroplast genome database was firstly constructed for flowering plants (4,161 from NCBI and 271 newly sequenced ones generated by our group) for KrakenUniq version 0.5.5 [86]. For gut metagenome data, we filtered out reads mapped to the *A. cerana* genome or to the *de novo* bacterial assemblies, and used the remaining reads for pollen diet profiling. The remaining reads were first aligned to the customized chloroplast genomes with KrakenUniq [86] with default parameters. Those mapped reads were aligned to nt database with blastn with e-value setting as 1e-5, and the best alignment were retained. Then the reads from the alignments with similarity > 95% and query coverage > 90% to reference sequences from plants were kept, and used to estimate the pollen abundance at the family level. The families with a relative abundance less than 1% were filtered out.

The geographical variation in pollen composition was also conducted with metagenome data from honey samples collected from five representative regions of this study (SC_AB, SC_GB, SX_QL, QH_GD, JL_DH, Table S11) [87]. For each sample, pollen pellets were centrifuged from diluted honey at 12,000 rpm for 15 min, and used for DNA extraction following Soares et al [88]. DNA samples were sequenced using Illumina HiSeq X Ten platform (350 bp insert size; 150 PE). Reads were assembled with MitoZ assembly module [89], with a K-mer size of 31 bp, a minimum edge depth of 3 and a minimum output length of 500 bp. The assemblies were aligned to the nt database with blastn and the best alignment was retained. Then the assemblies with similarity > 95% and query coverage > 90% to reference plant sequences were retained. Clean reads were then aligned to these assemblies using Minimap2 [60], and the mapped reads were used to estimate the pollen abundance at the family level with SamBamba [90]. The families with a relative abundance less than 1% were filtered out.

The gut bacterial phylotype and KO composition from *de novo* assembly and annotation were used in the correlation analysis with pollen composition at the family level.

### Heritability of bacterial diversity

The rank-based inverse normal transformation of the relative abundance with reference-based method was used in the heritability analysis. The heritability was defined as the Percentage of Variance Explained (PVE) and estimated with Genome-wide Ecient Mixed Model Association (GEMMA, v0.94) [91]. To control the effects of individual relatedness, population structure and diet variation, we regressed the transformed gut bacteria abundance with the first three PCs from the PCA of the host genotypic data, and the pollen Shannon index from the gut. Then PVE estimation was performed with the residuals using GEMMA (with relatedness matrices and the HE regression algorithm). A phylotype or SDP was considered heritable if the PVE measurements did not show overlaps with zero.

### The association between host genetic variation and bacterial diversity

The rank-based inverse normal transformation of the relative abundance of core gut bacteria was used in the Genome-Wide Association Studies (GWAS) analysis. We used the Linear Mixed Model in rMVP v1.0.0 [92]. In the GWAS analysis, the kinship between individuals, the first three PCs in host PCA, and the diet (Shannon index of pollen family composition) were used for correction. We used the ‘EMMA’ method to analyze variance components in rMVP. The statistical significance level was set to *P* < 5×10^-8^ for the GWAS association.

### The effects of diet on the abundance of *Gilliamella* and *Lactobacillus* Firm-5

A *Gilliamella* strain (B2889, belonging to the SDP Gillia_Acer_2) were cultivated with HIA, and a *Lactobacillus* Firm-5 strain (B4010) were cultivated with MRS with 0.02 g/ml D-frutcose (aladdin F108331) and 0.001 g/ml L-cysteine (aladdin C108238). The microbiota-free *A. cerana* workers were obtained following Zhang *et al*. [80]. Pupae in late stage were removed from brood frames and incubated in sterile plastic bins at 35 °C. Both bacterial strains of OD_600_=1 were mixed at equal volumes, and then mixed with 50% (v/v) sterilized sucrose syrup, which were fed to newly emerged microbiota-free honeybees. After three days, cellobiose (Shanghai Yuanye Bio-Technology Co., Ltd S11030, final concentration 5 mg/mL) and solutions with different proportions of pectin (Sigma, P9135) and cellulose (Megazyme, P-CMC4M) (1:10, 10:1, final mixed concentration 5.5 mg/mL) mixed with sterilized 50% sucrose syrup were fed to honeybees, respectively. Honeybees fed with only 50% sucrose syrup were used as control. After feeding for four days, DNA were extracted from bee guts and used for the qPCR assay.

### qPCR assay

We conducted real-time qPCR experiments to determine bacterial loads for both *Gilliamella* and *Lactobacillus* Firm-5 after the feeding experiments. 16S-F-Gillia (5’-TGAGTGCTTGCACTTGATGACG-3’) and 16S-R-Gilla (5’-ATATGGGTTCATCAAATGGCGCA-3’) primers were used for *Gilliamella* 16S rRNA gene amplification. 16S-F-Firm5 (5’-GCAACCTGCCCTWTAGCTTG-3’) and 16S-R-Firm5 (5’-GCCCATCCTKTAGTGACAGC-3’) primers [93] were used for *Lactobacillus* Firm-5 16S rRNA gene amplification. Actin-AC-F (5’-ATGCCAACACTGTCCTTTCT-3’) and Actin-AC-R (5’-GACCCACCAATCCATACGGA-3’) primers were used to amplified *actin* gene of the host *A. cerana* [94], which was used to normalize the bacterial amplicons [93]. The 16S target sequences were cloned into vector pEASY-T1 (Transgen) and the Actin target sequence was cloned into pCE2 TA/Blunt-Zero Vector (Vazyme), then confirmed by Sanger sequencing. The copy number of the plasmid was calculated, serially diluted and used as the standard. qPCR was performed using the ChamQ Universal SYBR qPCR Master Mix (Vazyme) and QuantStudio 1 (Thermo Fisher) in a standard 96-well block (20-μL reactions; incubation at 95 °C for 3 min, 40 cycles of denaturation at 95 °C for 10 s, annealing/extension at 60 °C for 20 s). The data were analyzed using the QuantStudio Design & Analysis Software v1.5.1 (Thermo Fisher) and Excel (Microsoft). *P* values were calculated using the Mann-Whitney test.

## List of Abbreviations

gANI: genome-wide average nucleotide identity
SDP: sequence-discrete population
SNV: single nucleotide variation
CNV: copy number variation
GH: glycoside hydrolase
PL: polysaccharide lyase
PTS: phosphotransferase system
ABC: ATP binding cassette
CGC: CAZyme gene cluster
PGA: polygalacturonic acid
HIA: heart infusion agar
MRS: De Man, Rogosa and Sharpe
TPY: trypticase-phytone-yeast
PVE: Percentage of Variance Explained
GEMMA: Genome-wide Ecient Mixed Model Association
PCoA: principal coordinate axis
GWAS: Genome-Wide Association Studies

## Declarations

## Acknowledgements

Many colleagues and collaborators have facilitated sampling or contributed samples to this work, including Kang Lai, Yongkun Ji, Jieluan Li, Dandan Lang, Shuang Yang, Shijie Liang, Zhanhai Cai, Dingqian Xiang, Jie Deng, Hua Liang, Kangkai Zheng, Abuduoji, Liang Li, Jihuan Du, Zijing Zhang, Xuhui Wang, Zhaohuai Li, Feixiang Wang, Yanqiong Peng, Jialong Huang and Zhengwei Wang. Chenyi Li, Yating Du, Chengfeng Yang, Lizhen Guo, Lifei Qiu and Wenjun Zhang from China Agricultural University isolated and cultured honeybee gut bacteria that were used for genome sequencing in this work. We thank Chengfeng Yang from China Agricultural University for providing the primers and plasmid of *Gilliamella* used for qPCR assay. We thank Prof. Hao Zheng from China Agricultural University for his comments on early drafts of the manuscript.

## Authors’ contributions

Xin Z. designed the study. Xin Z., S.L. and Xue Z. organized and coordinated the study. Q.S. coordinated sample collection, bacterial isolation and genome annotation, reference-based metagenome mapping, and SDP analysis. M.T. conducted *de novo* assembly of bacteria and metagenome, and metagenomics analysis. S.L. conducted genetic variation analysis of *A. cerana*, heritability and GWAS analysis. J.H. conducted diet profiling. Q.S. and J.T. conducted feeding experiments and qPCR assay. Xin Z., Q.N. and X.L. organized sampling. S.L, Xin Z. and Xuguo Z. wrote the first drafts and all authors contributed to and proofed the manuscript.

## Availability of data and materials

Raw data for metagenome and bacteria strain genome have been deposited in the bio-project PRJNA705951 in NCBI. Raw honey sequencing data were deposited in the bio-project PRJNA722869 in NCBI. In-house scripts are available from the corresponding authors upon request.

## Ethics approval and consent to participate

Not applicable.

## Consent for publication

Not applicable.

