## Supplemental figures and tables for "Diet outweighs genetics in shaping gut microbiomes in Asian honeybee": SOM_20211108.docx

**Supplementary Figures**

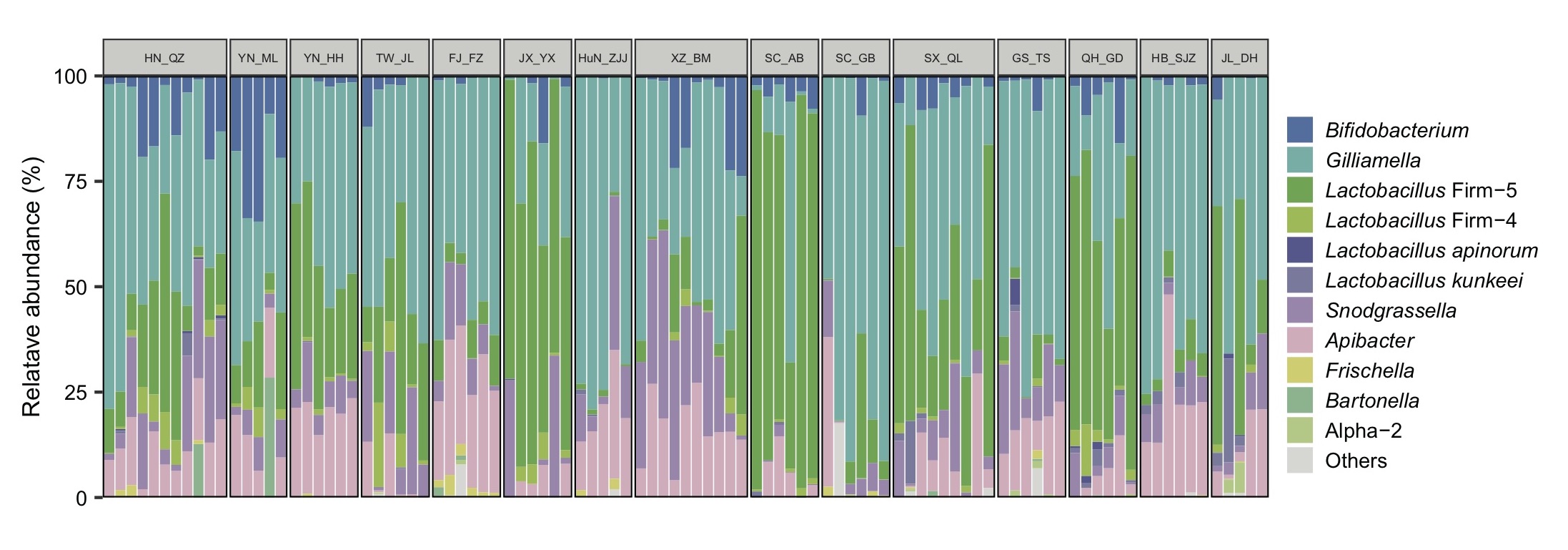

**Figure S1. Community composition and diversity at the phylotype level in gut microbiota from *A. cerana*.** The results were analyzed by Kraken2 with 390 bee gut bacteria genomes as reference in Supplementary Table 2. Host populations are indicated by short bars at the top.

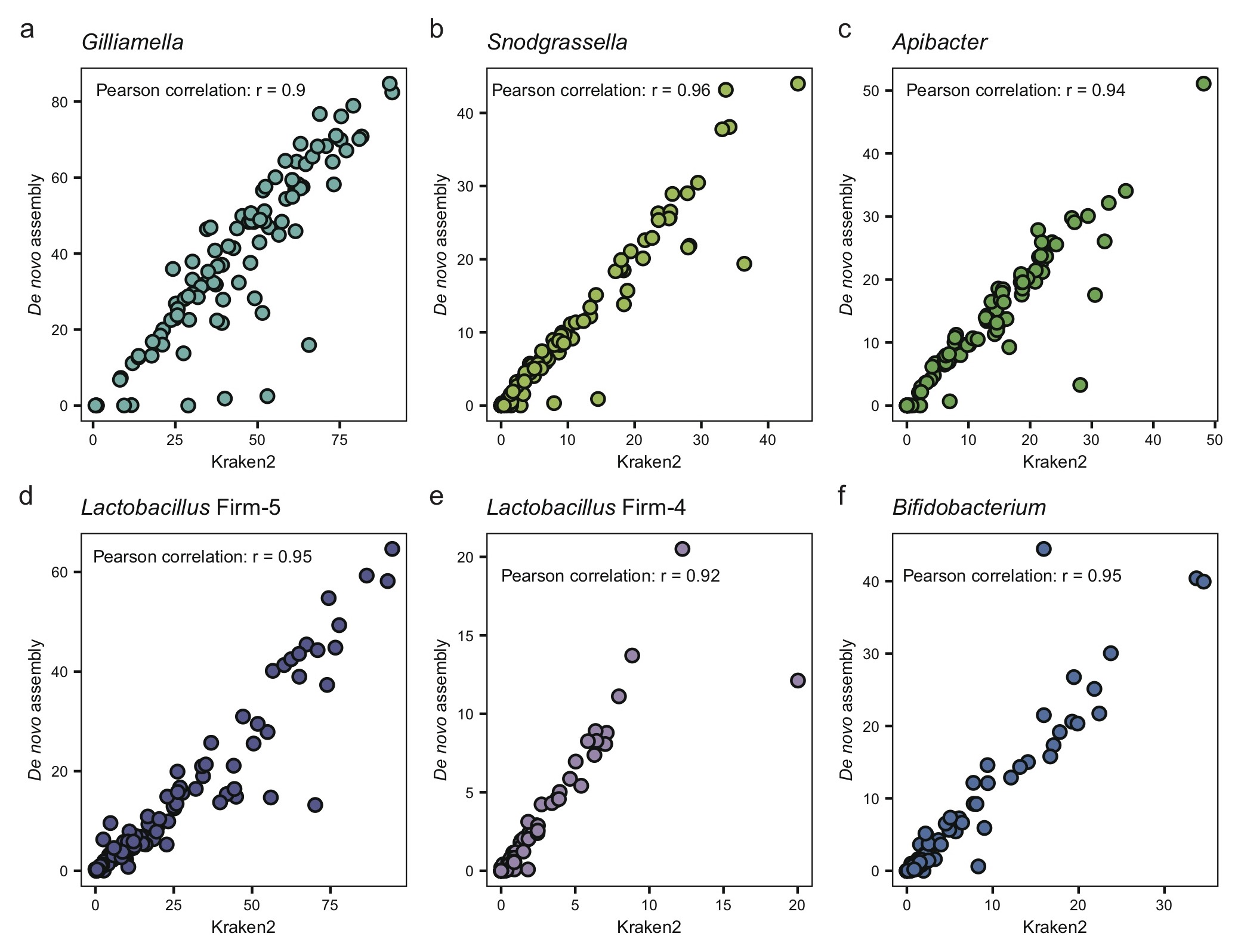

**Figure S2. The relative abundance of core bacteria with the *de novo* methods were highly correlated with the results from reference-based Kraken2 methods.**

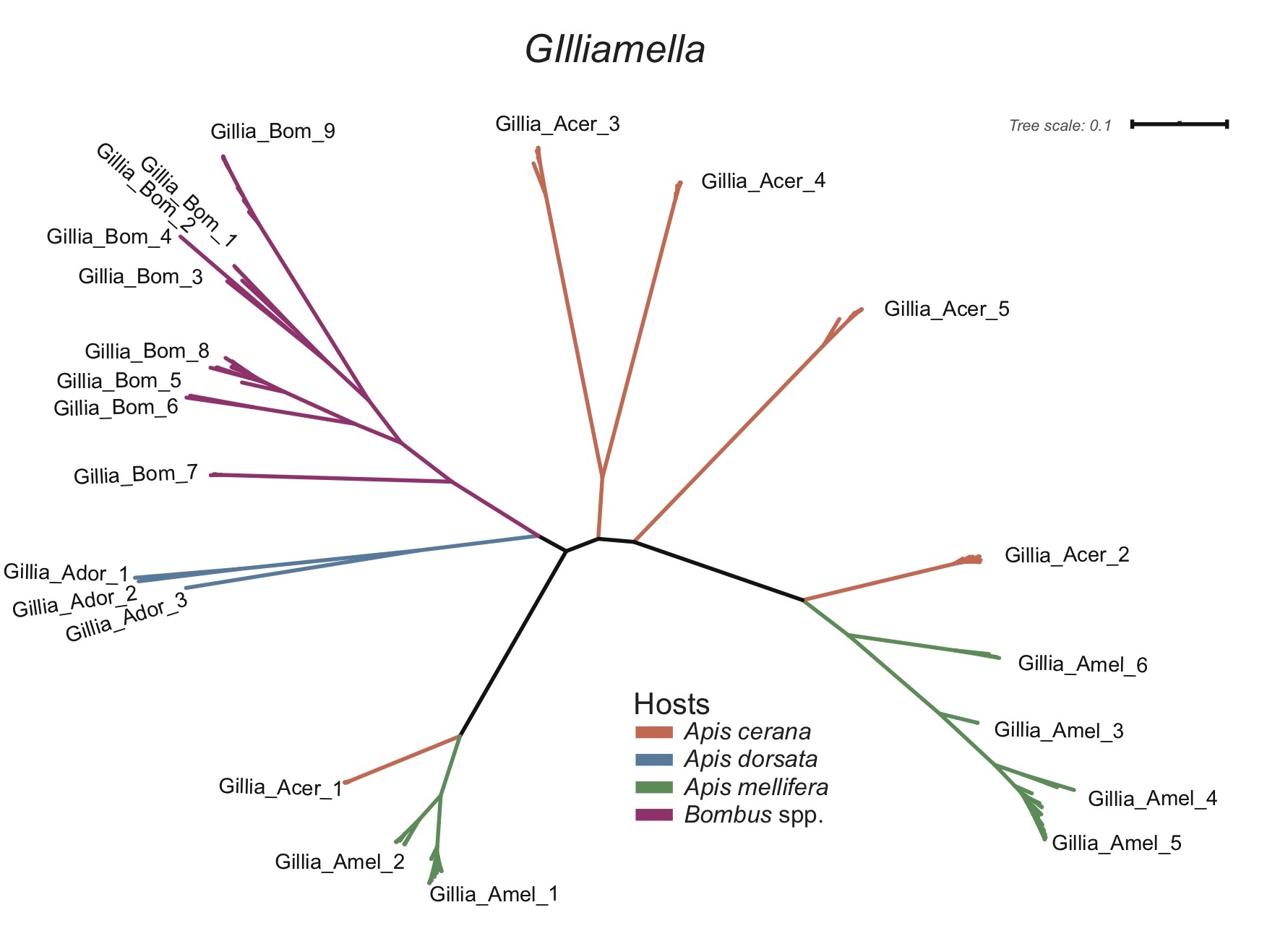

**Figure S3. Maximal-likelihood (ML) phylogenetic tree of *Gilliamella* strains from bees.** ML tree was built on the core genes existing in all strain genomes. Branch colors correspond to host species.

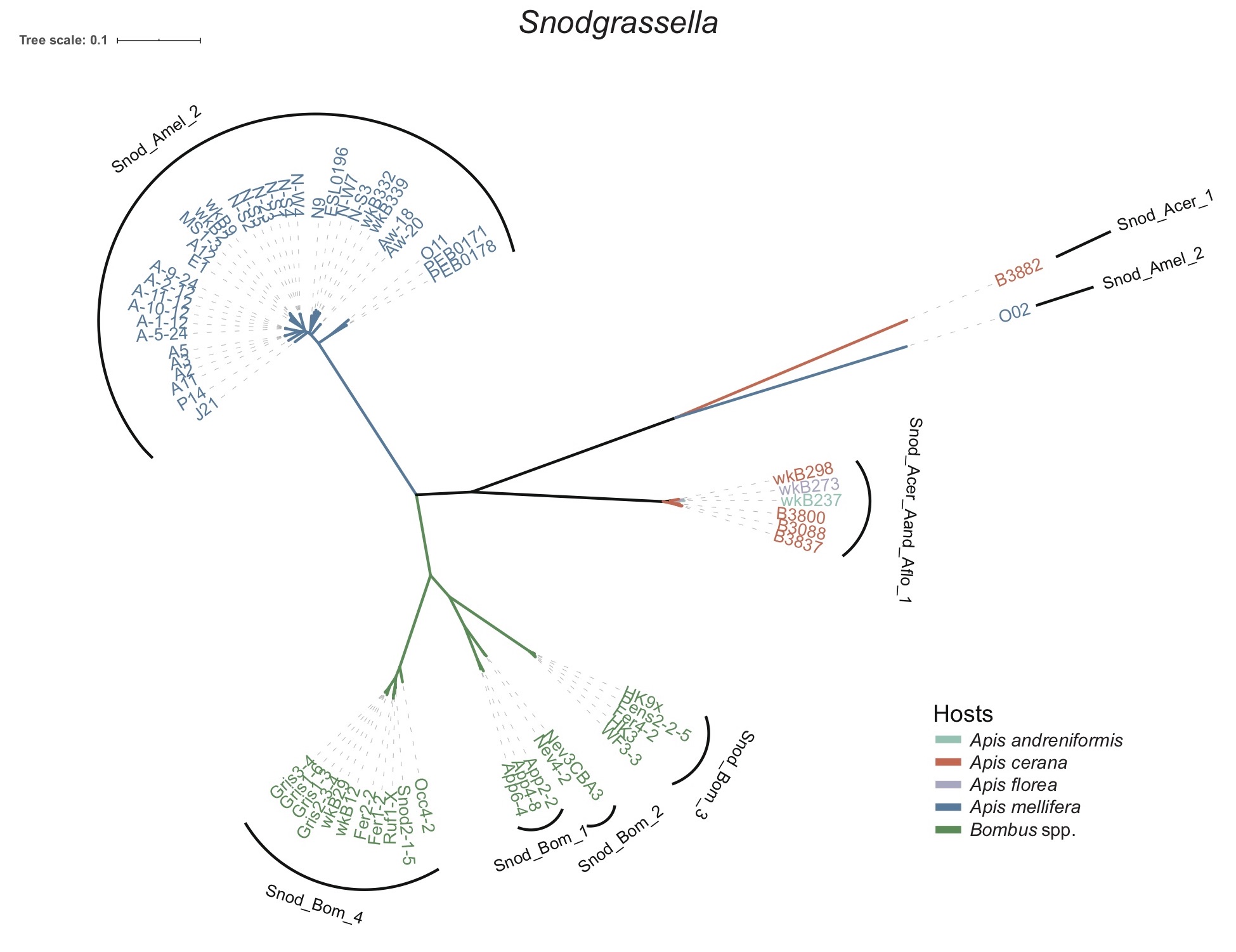

**Figure S4. Maximal-likelihood (ML) phylogenetic tree of *Snodgrassella* strains from bees.** ML tree was built on the core genes existing in all strain genomes. Branch colors correspond to host species.

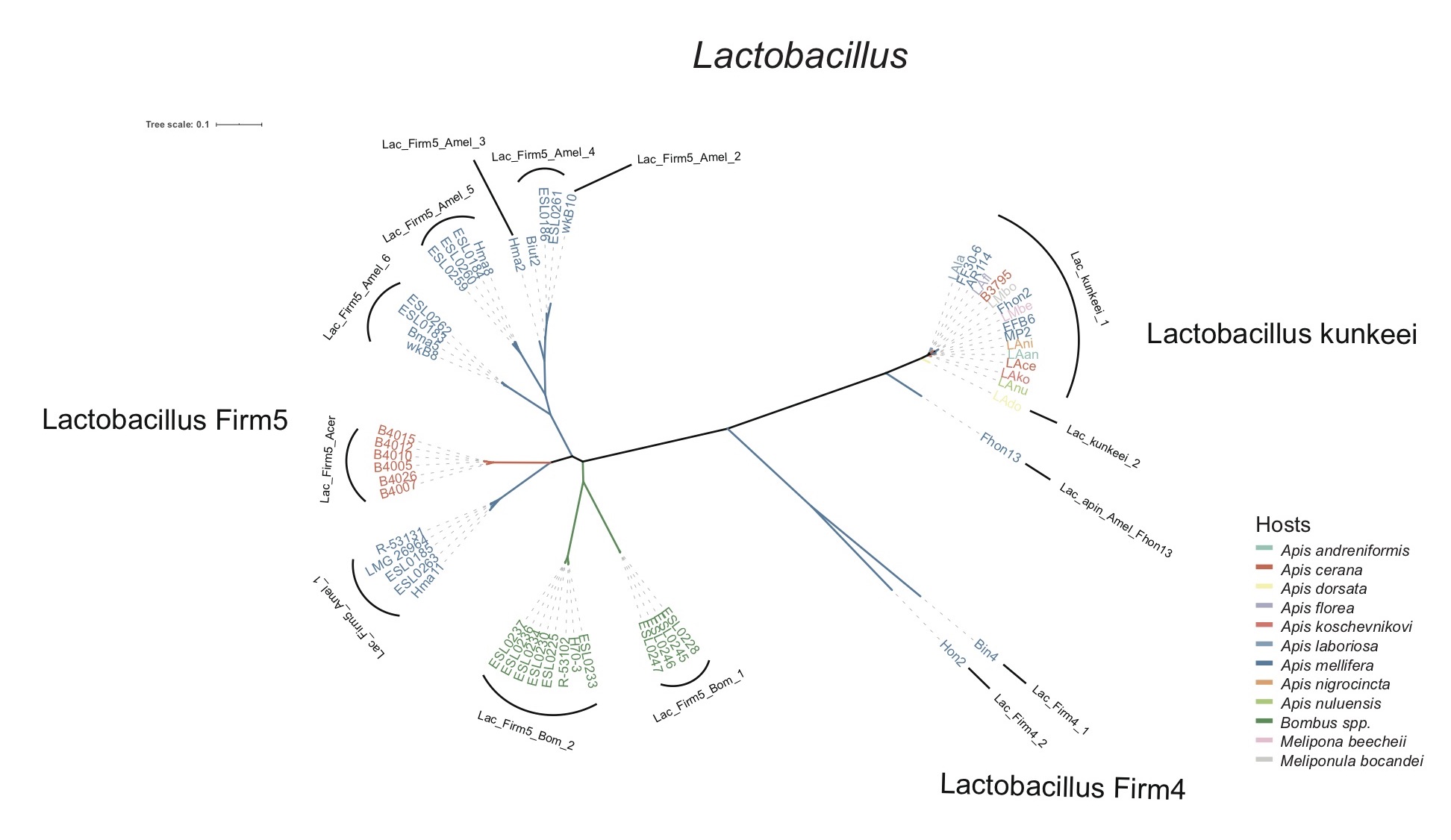

**Figure S5. Maximal-likelihood (ML) phylogenetic tree of *Lactobacillus* strains from bees.** ML tree was built on the core genes existing in all strain genomes. Branch colors correspond to host species.

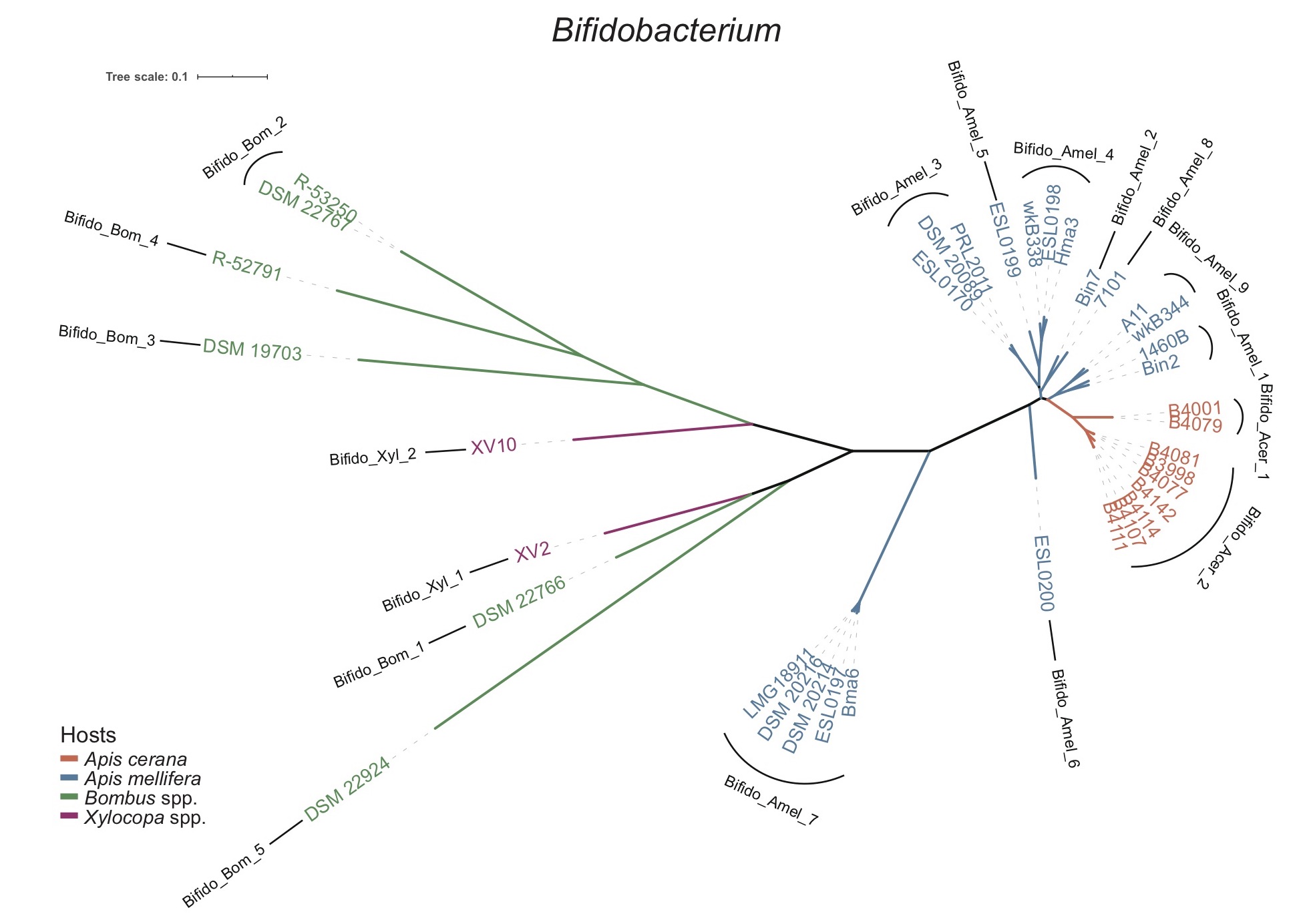

**Figure S6. Maximal-likelihood (ML) phylogenetic tree of *Bifidobacterium* strains from bees.** ML tree was built on the core genes existing in all strain genomes. Branch colors correspond to host species.

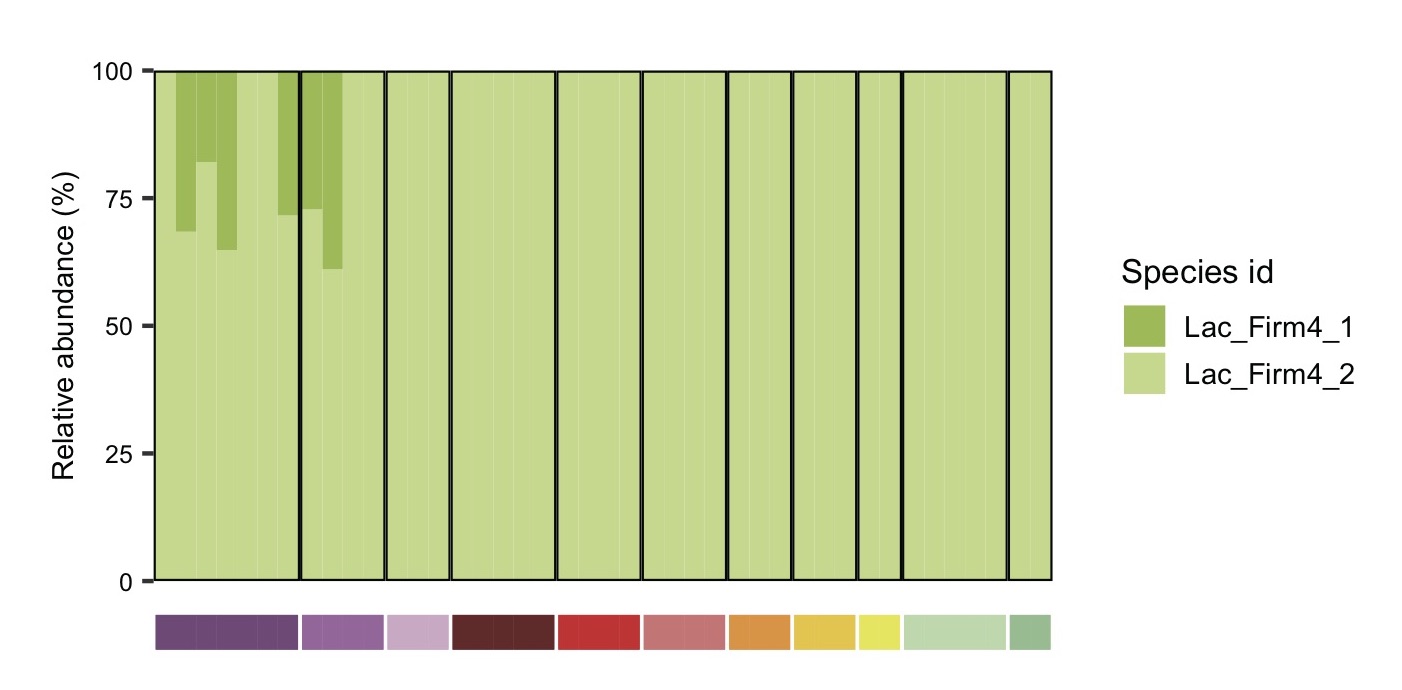

**Figure S7. The SDP composition of *Lactobacillus* Firm-4 in gut metagenome samples of *A. cerana* estimated by MIDAS.**

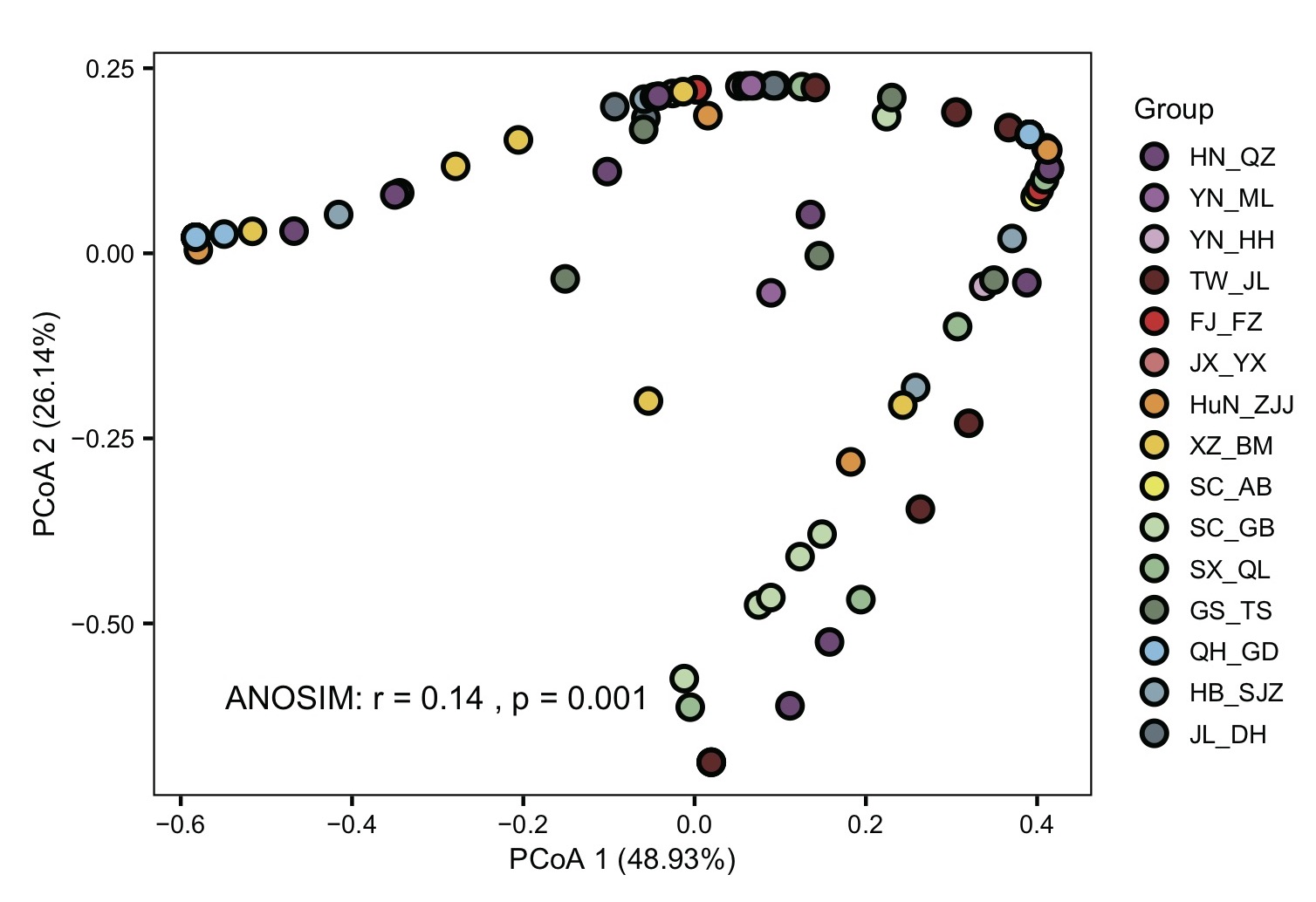

**Figure S8. PCoA of *Gilliamella* SDP composition in populations of *A. cerana*.**

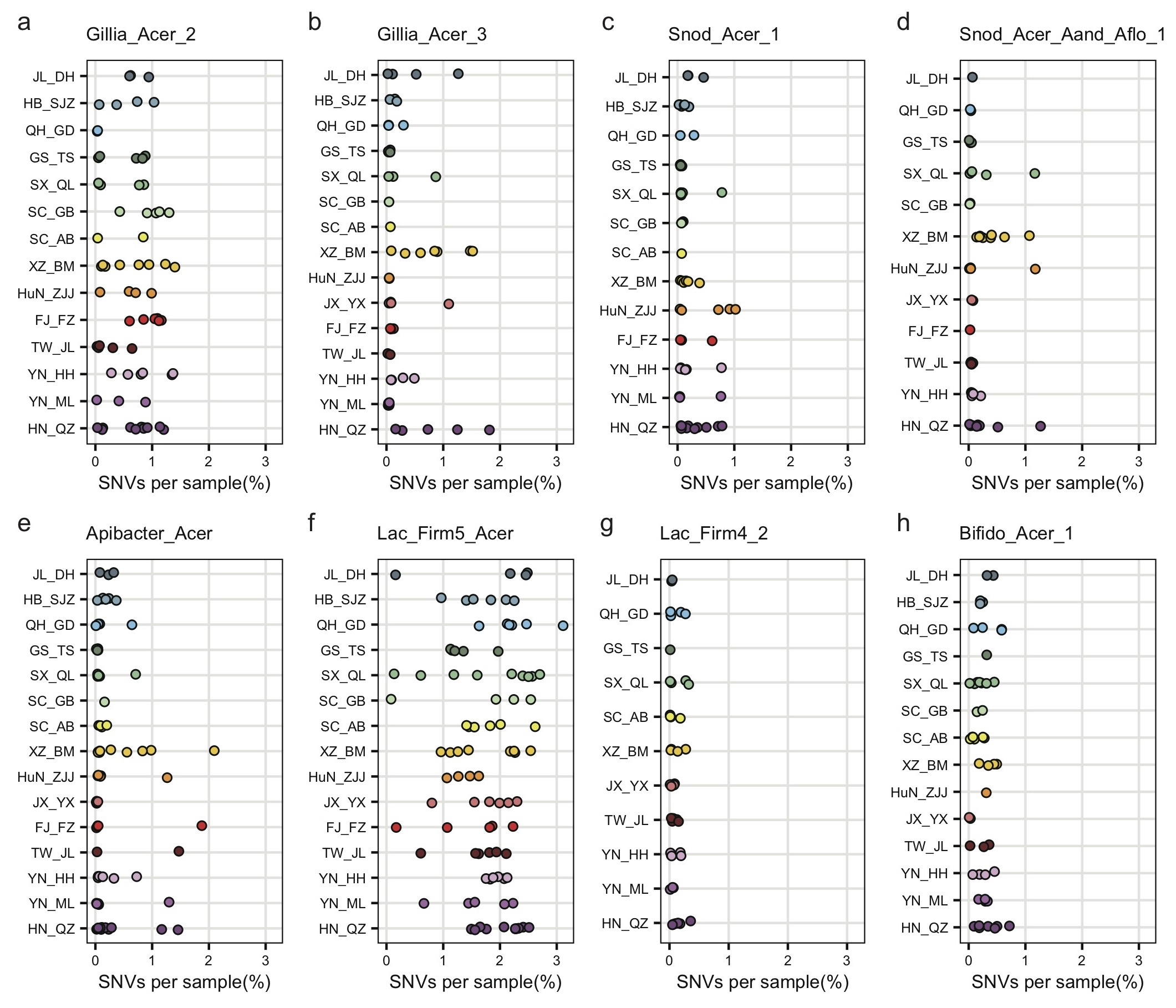

**Figure S9. Fraction of SNVs for the dominant SDPs varied in populations of *A. cerana*.** Dominant SDPs include those show relatively high frequency in all samples. SNVs in each sample were calculated from all SNPs existing in each SDP.

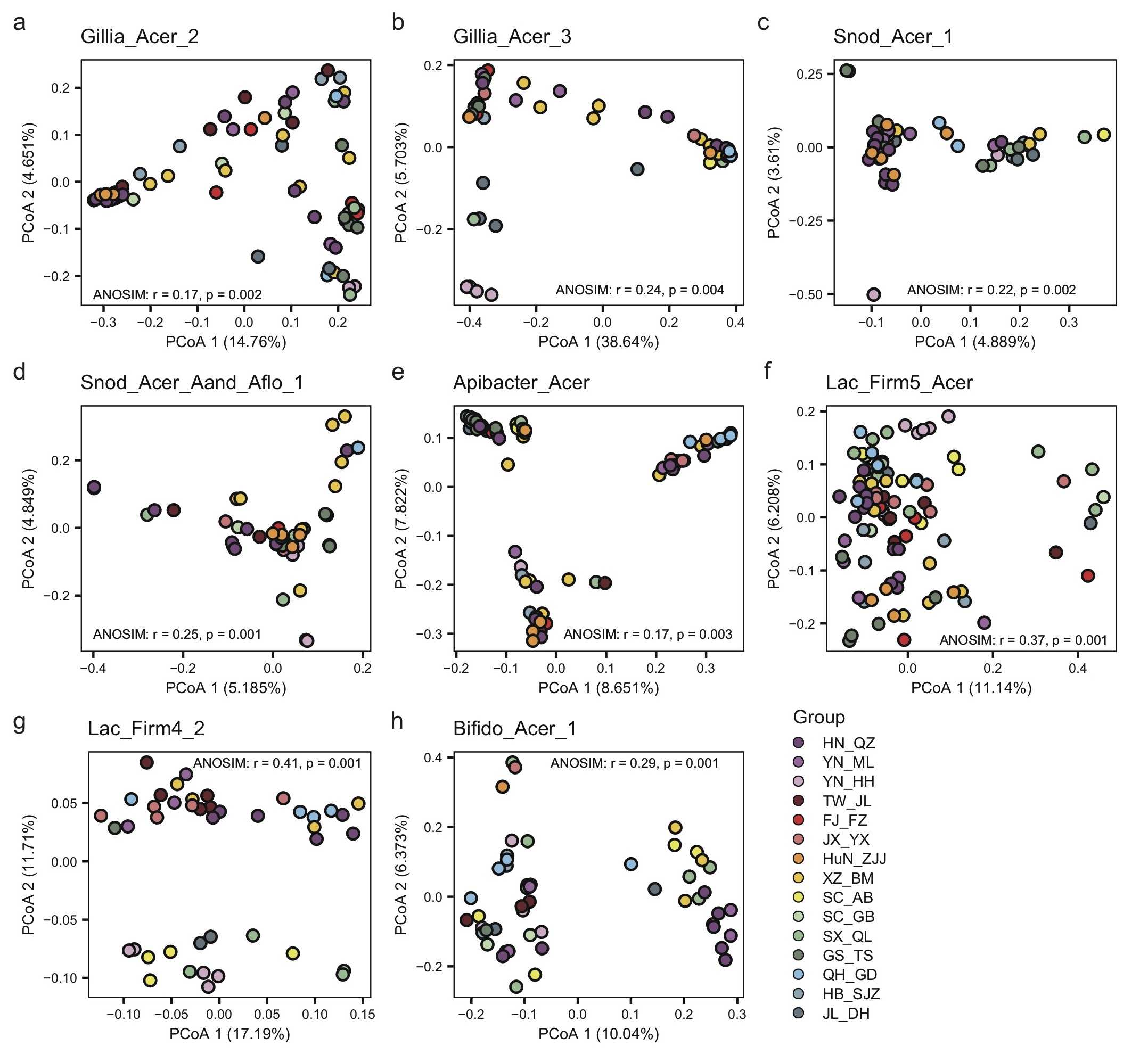

**Figure S10. SNV differences for dominant SDPs in the gut microbiome from 15 populations of *A. cerana*.**

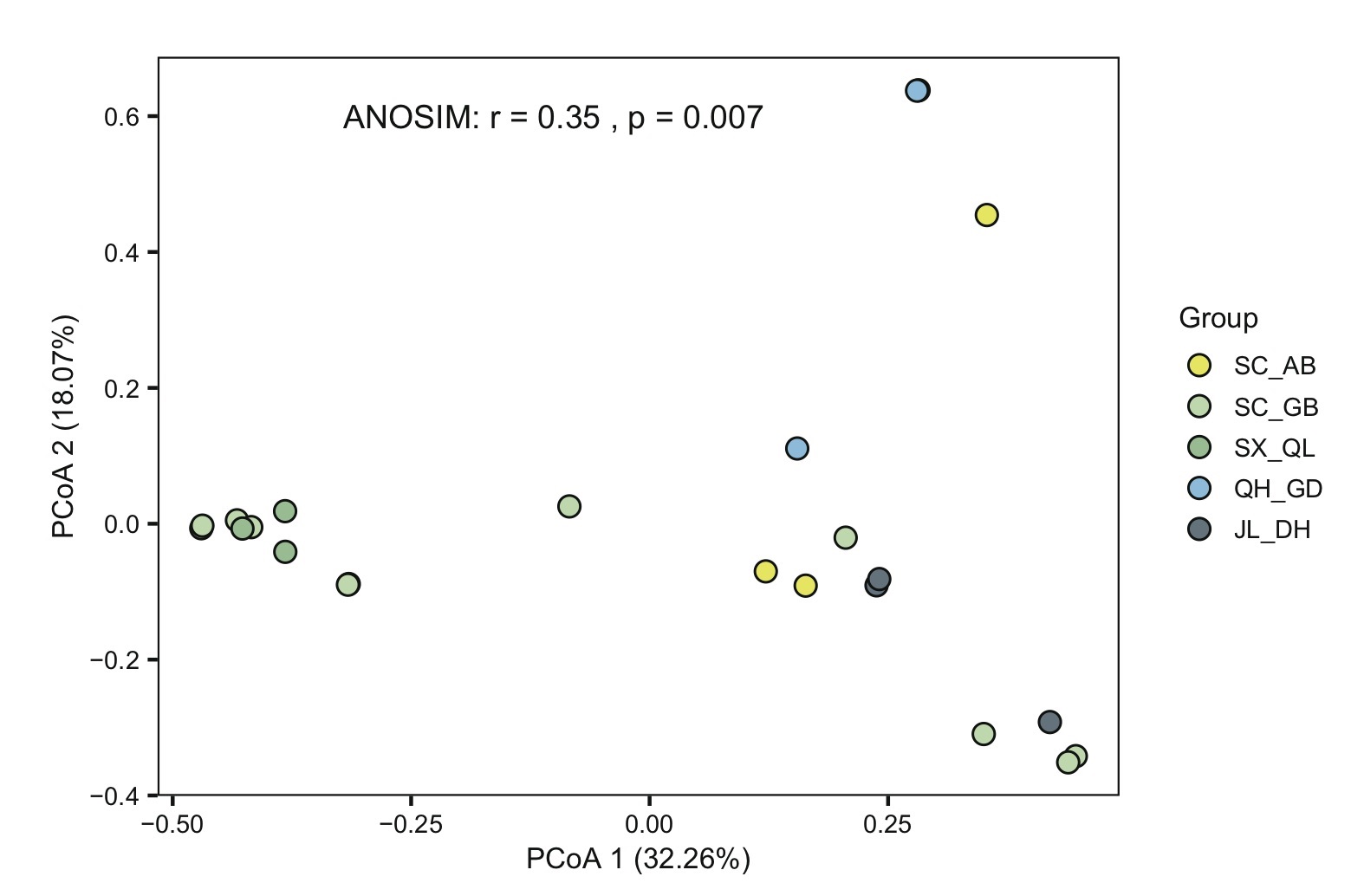

**Figure S11. The pollen composition at the family level varied in honey from populations of *A. cerana*.**

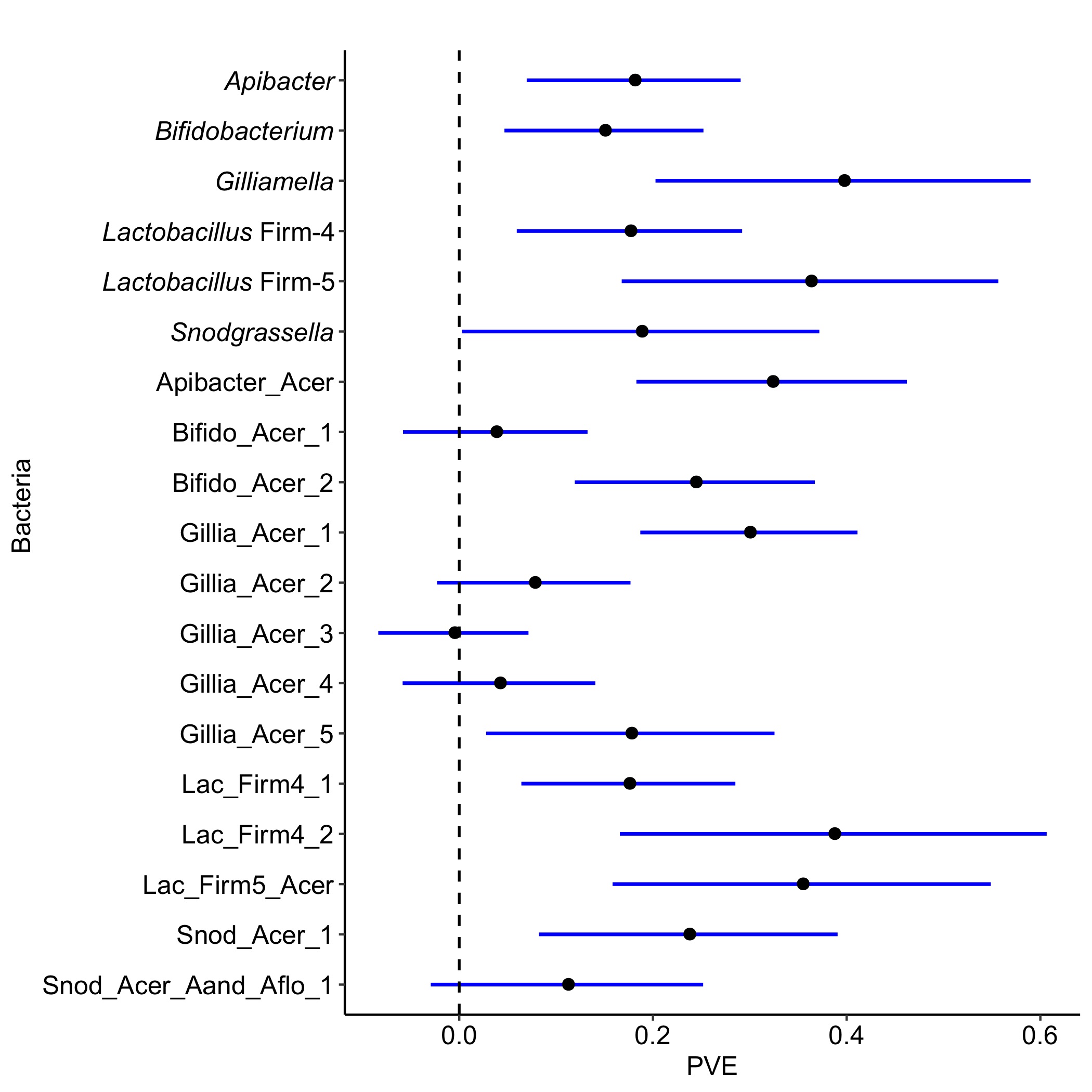

**Figure S12. SNP heritability of the core phylotypes and SDPs for gut microbiota in *A. cerana*.** Each point represents the estimated PVE with GEMMA by genome-wide SNPs for the abundance of the core phylotypes and SDPs. Bars indicate SE measurements around the estimate. PVE: Percentage of Variance Explained.

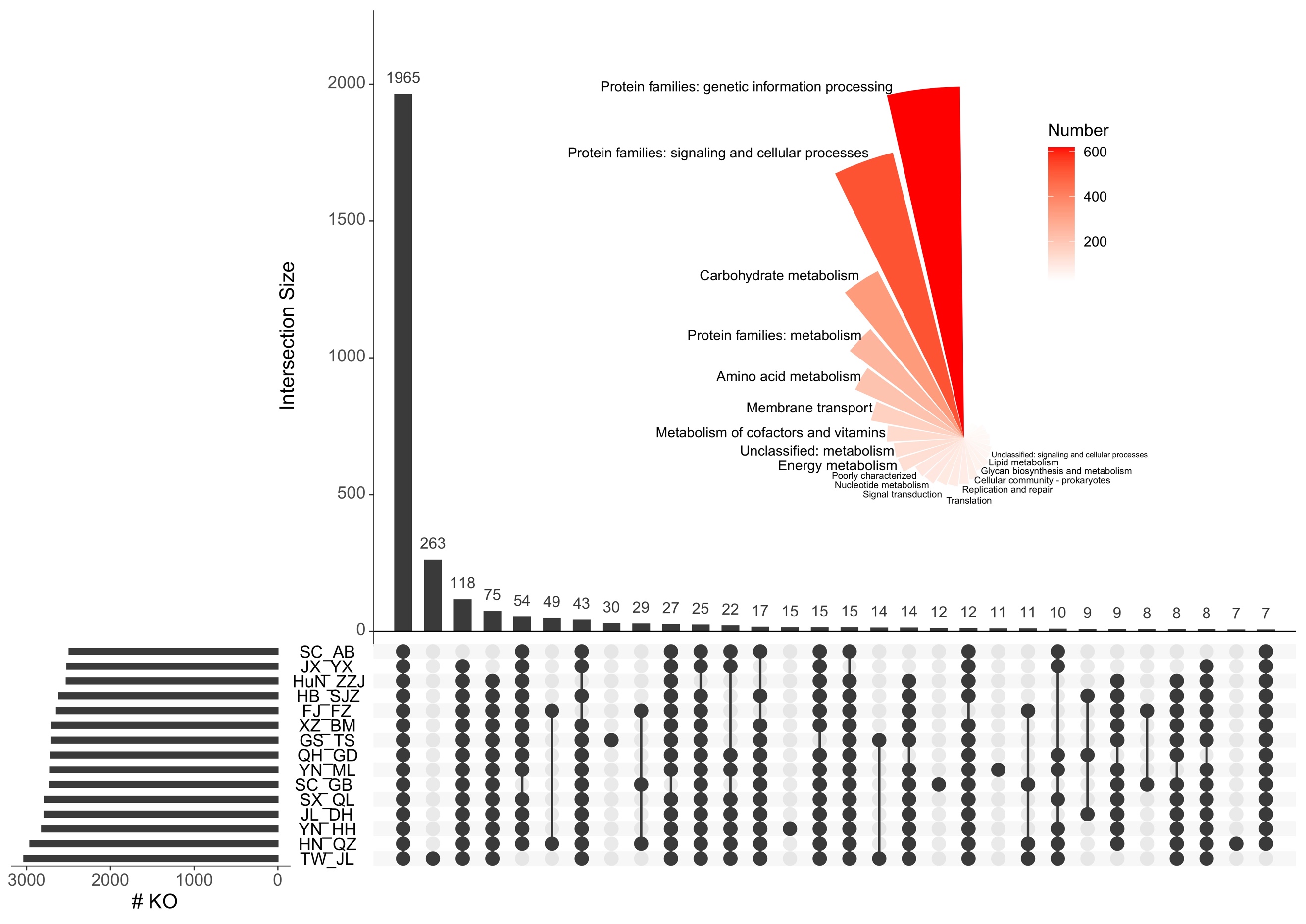

**Figure S13. Distribution of KEGG orthologs (KOs) in gut microbes from 15 populations of *A. cerana*.** Horizontal bars on the bottom left represent KO numbers annotated in each population. Vertical bars indicate the numbers of KOs exclusively associated with one or multiple groups. The common KO numbers in each category are in red. The area of each KO category is proportional to the KO number.

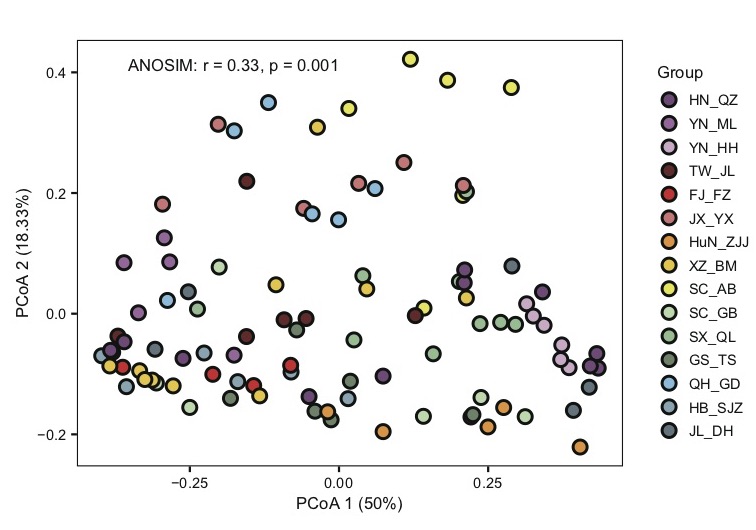

**Figure S14. Gut microbe KO abundance showed significant difference among populations with PCoA.**

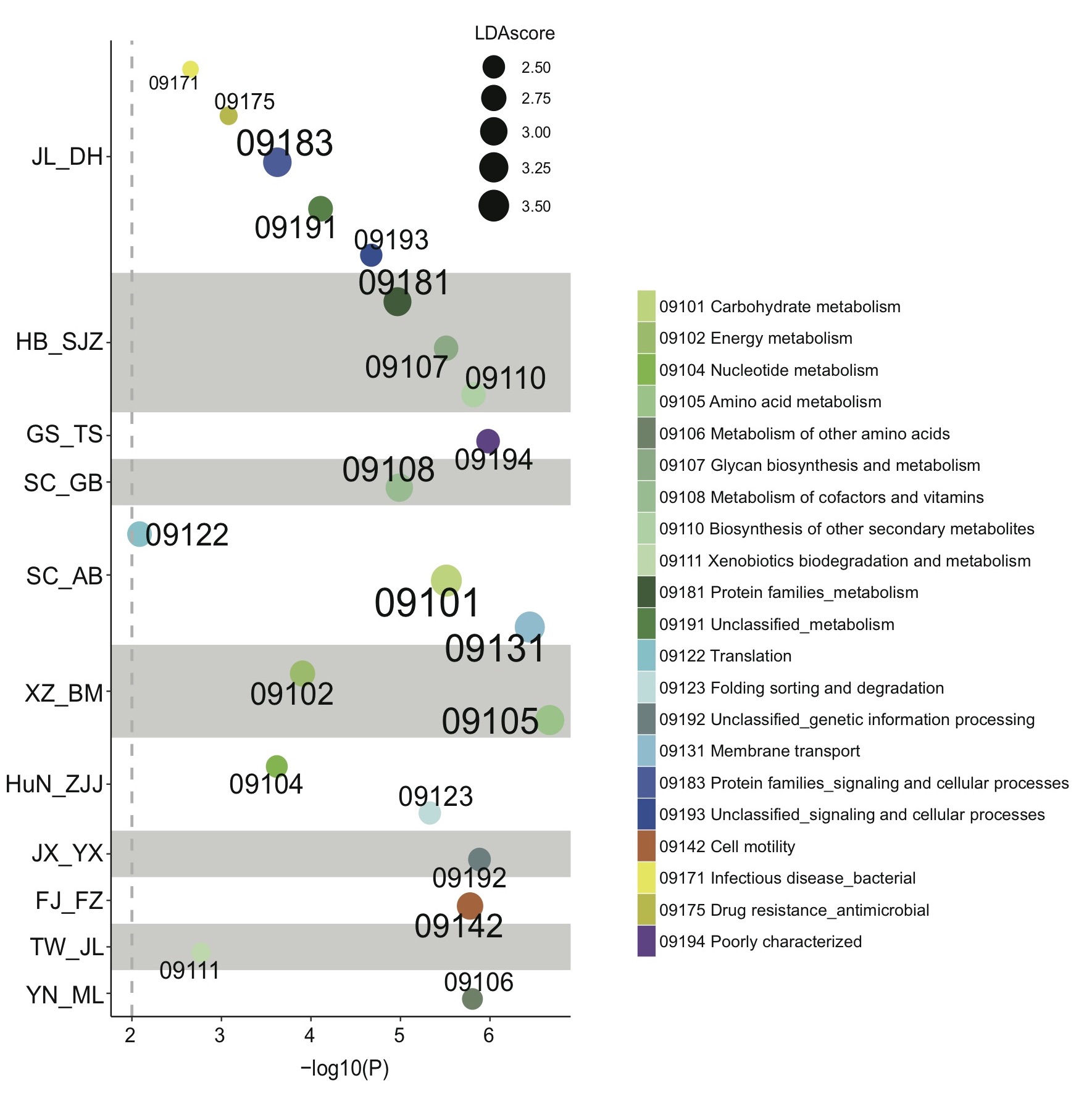

**Figure S15. Characterized KEGG categories in gut microbiota from populations of *A. cerena* using LEfSe algorithm.** The size of the squares represents the LDA score.

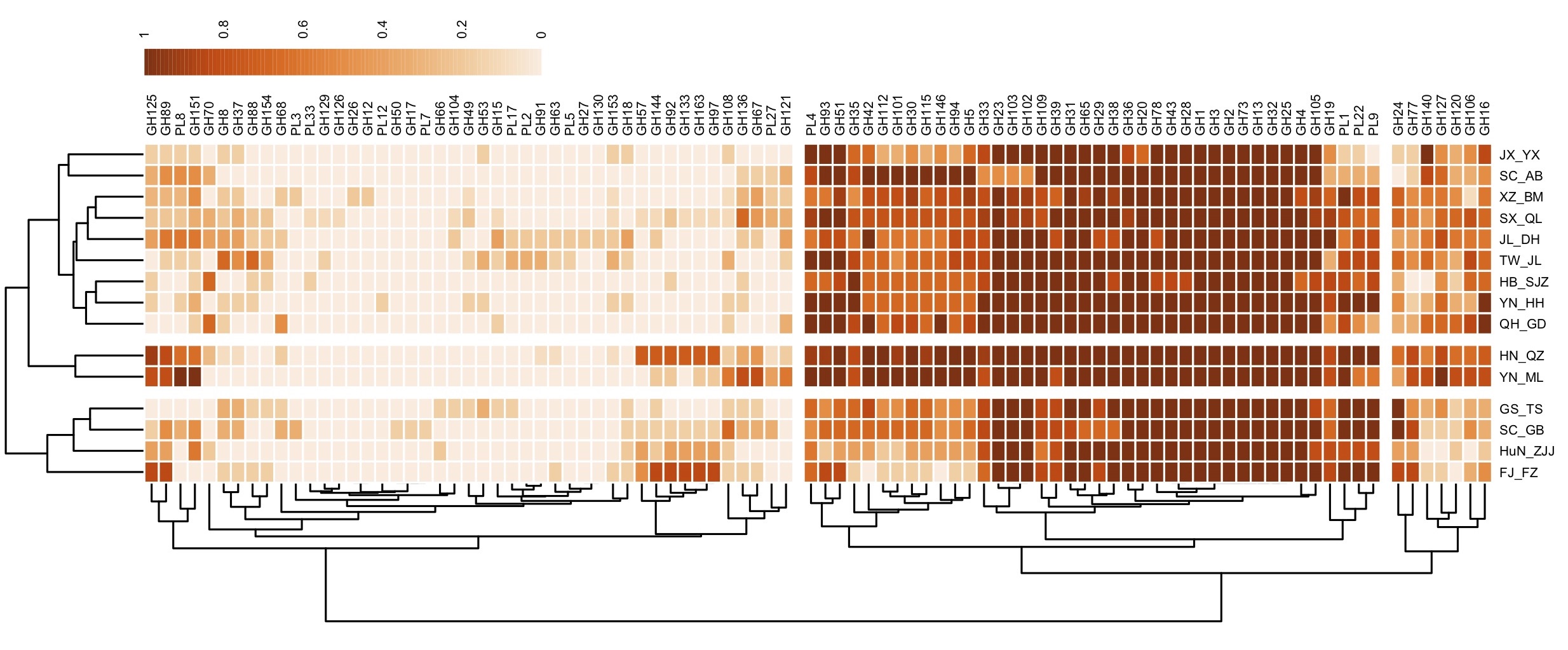

**Figure S16.** **Glycoside hydrolases (GH) and polysaccharide lyases (PL) gene profiles in gut microbiota from populations of *A. cerana.*** Color represents the frequency of the GH/PL family in each population.

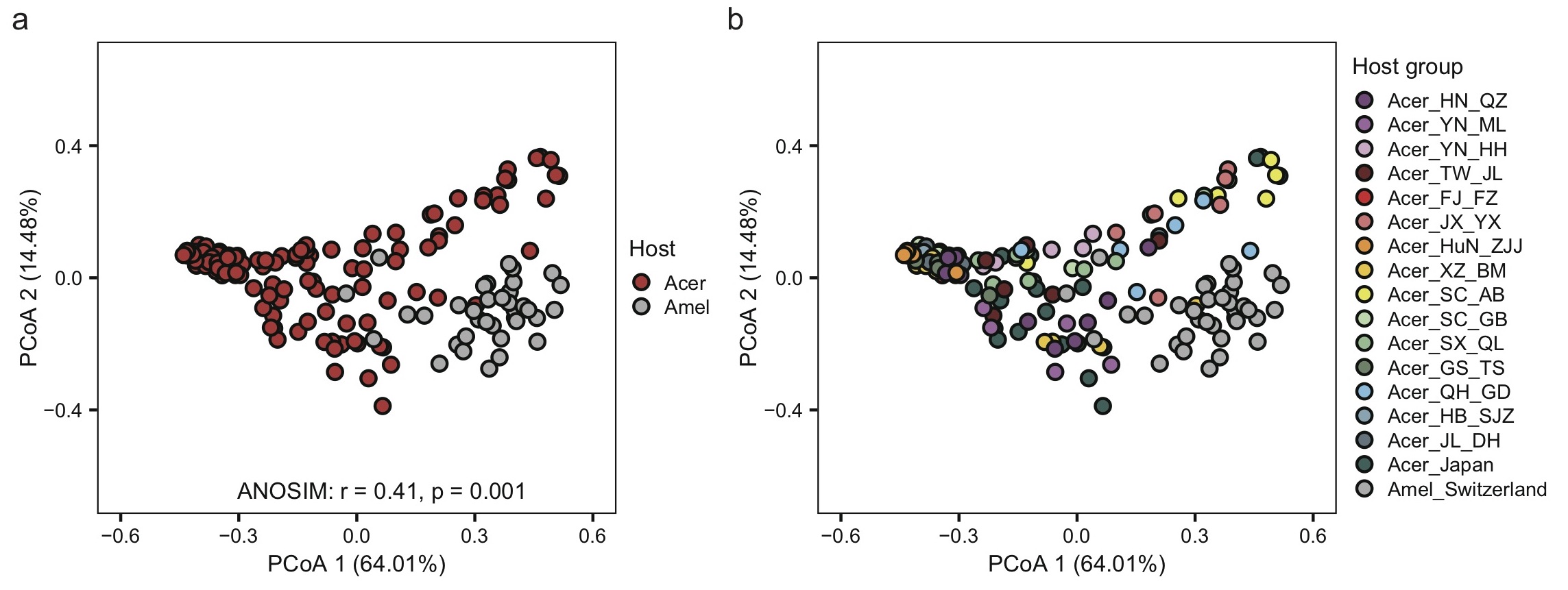

**Figure S17. Gut microbe community composition showed significant difference between *A. cerana* and *A. mellifera* with PCoA.** (A) Gut microbe community difference at the host species level. (B) Gut microbe community composition showed variation among different populations of *A. cerana* (including population from Japan) with PCoA.

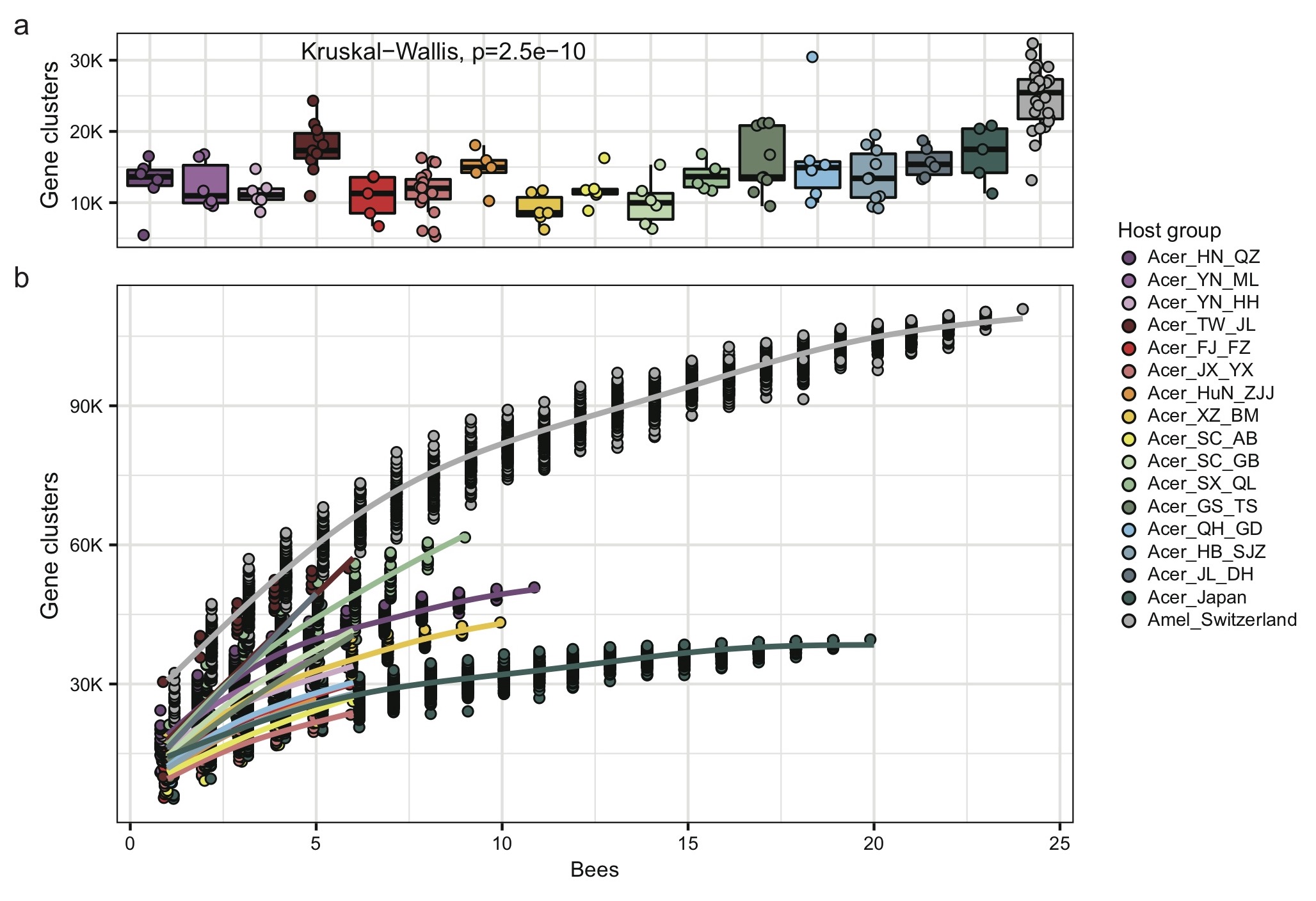

**Figure S18. Gene cluster numbers in gut microbiota varies in *A. cerana* and *A. mellifera*.** All comparisons were based on the 400Mb bacteria-mapped reads. (a) Gene cluster numbers per sample showed difference in different populations of *A. cerana* and *A. mellifera*. (b) Cumulative number for gene cluster numbers in gut metagenome from *A. mellifera* and each population of *A. cerana*.
